# Neuronal HLH-30/TFEB modulates muscle mitochondrial fragmentation to improve thermoresistance in *C. elegans*

**DOI:** 10.1101/2022.04.07.487519

**Authors:** Shi Quan Wong, Catherine J Ryan, Louis R Lapierre

## Abstract

Transcription factor EB (TFEB) is a conserved master transcriptional activator of autophagy and lysosomal genes that modulates organismal lifespan regulation and stress resistance. As neurons can coordinate organism-wide mechanisms, we investigated the role of neuronal TFEB in stress resistance and longevity. To this end, the *C. elegans TFEB* orthologue, *hlh-30*, was rescued panneuronally in *hlh-30* loss of function mutants. While important in the long lifespan of *daf-2* animals, neuronal *hlh-30* was not sufficient to restore normal lifespan in short-lived *hlh-30* mutants. However, neuronal HLH-30/TFEB rescue mediated robust improvements in the heat stress resistance of wild-type but not *daf-2* animals. Notably, these mechanisms can be uncoupled, as neuronal HLH-30/TFEB regulates longevity and thermoresistance dependently and independently of DAF-16/FOXO respectively. Through transcriptomics profiling and functional analysis, we identified the uncharacterized gene *W06A11.1* as a bona fide mediator of heat stress resistance via the induction of mitochondrial fragmentation in distal muscles. Neuron-to-muscle communication occurred through a modulation of neurotransmission. Taken together, this study uncovers a novel mechanism of heat stress protection mediated by neuronal HLH-30/TFEB.

## INTRODUCTION

Transcription factor EB (TFEB) transcriptionally induces genes of the autophagy and lysosomal pathway, a process that recycles cellular components through lysosomal degradation (Settembre *et al.*, 2011). The transcriptional activity of TFEB is dependent on its nuclear translocation, which is modulated through phosphorylation by mammalian target of rapamycin complex 1 (mTORC1), dephosphorylation by calcineurin, and nuclear export by exportin 1 (Silvestrini *et al.*, 2018; Napolitano *et al.*, 2018; Li *et al.*, 2018; Martina *et al.*, 2012; Roczniak-Ferguson *et al.*, 2012; Settembre *et al.*, 2012; Medina *et al.*, 2015). In addition to its role as an autophagy and lysosomal inducer, studies in *C. elegans* have uncovered the diverse organismal processes regulated by the TFEB ortholog, HLH-30, including longevity (Lapierre *et al.*, 2013; Lin *et al.*, 2018), adult reproductive diapause (Gerisch *et al.*, 2020), resistances to heat and oxidative stresses (Lin *et al.*, 2018), starvation (O’Rourke and Ruvkun, 2013; Harvald *et al.*, 2017; Settembre *et al.*, 2013), and pathogenic infection (Visvikis *et al.*, 2014; El-Houjeiri *et al.*, 2019; Wani *et al.*, 2021). Notably, HLH-30/TFEB mobilizes different transcriptional programs to regulate these processes, demonstrating its versatile role. How HLH-30/TFEB orchestrates these systemic responses from and between cell and tissue types remain unresolved and important to ascertain, as such signaling mechanisms are potential intervention targets for modulating aging and stress resistance.

Studies in *C. elegans* have revealed the important and conserved role of the nervous system in integrating stress signals and responses to modulate lifespan extension and organismal stress resistance in a non-cell autonomous manner (reviewed in (Miller *et al.*, 2020). In the regulation of longevity for instance, the rescue of DAF-16 forkhead Box O (FOXO) transcription factor in neurons partially regulated non-neuronal DAF-16/FOXO activity and lifespan extension in insulin/insulin-like growth factor receptor signaling (IIS)-defective *daf-2* mutants (Libina, Berman and Kenyon, 2003), whereas restoring IIS specifically to neurons abrogated the longevity phenotype of these mutants (Wolkow *et al.*, 2000). In the germlineless *glp-1* longevity model, lifespan extension mediated by microRNA *mir-71* in neurons required non-cell autonomous intestinal DAF-16/FOXO activity (Boulias and Horvitz, 2012). Neurons are also important for mediating the antagonistic activities of the nutrient sensors TORC1 and AMP-activated protein kinase on lifespan through the non-cell autonomous regulation of mitochondrial dynamics by neuropeptides (Zhang *et al.*, 2019).

In other *C. elegans* studies, the coordination of organismal stress responses by neurons have additionally been shown to be coupled to lifespan extension. For instance, constitutive neuronal endoplasmic reticulum (ER) unfolded protein response (UPR^ER^) activation by X-box binding protein 1 neuronal overexpression improved ER stress resistance and proteostasis, and extended lifespan through peripheral intestinal induction of the UPR^ER^, lysosomal activity, and lipid remodeling (Taylor and Dillin, 2013; Imanikia *et al.*, 2019a; Imanikia *et al.*, 2019b). In response to heat stress, neurons release neurotransmitters to other tissues to globally induce the heat shock response (HSR) (Prahlad, Cornelius and Morimoto, 2008). Furthermore, neuronal overexpression of the canonical HSR transcription factor heat shock factor 1 (HSF-1) not only enabled serotonin-mediated non-cell autonomous HSR induction in the absence of heat (Tatum *et al.*, 2015), but additionally conferred resistance to heat stress and lifespan extension (Douglas *et al.*, 2015). When faced with proteotoxic stress, neurons were capable of stimulating proteostasis distally by increasing molecular chaperone production in muscles through a non-cell autonomous signaling event termed transcellular chaperone signaling (O’Brien *et al.*, 2018). Overall, these findings highlighted that neurons constitute an essential site for lifespan modulation and organismal stress responses through non-cell autonomous mechanisms, although the exact intercellular and inter-tissue signaling events are largely undetermined.

Since HLH-30/TFEB is necessary for most longevity models (Lapierre *et al.*, 2013) and neurons are crucial in regulating longevity and stress resistance (Miller *et al.*, 2020), HLH-30/TFEB activity in neurons may be important in modulating these organismal phenotypes. Using *C. elegans*, we discovered that neuronal HLH-30/TFEB regulates both IIS-dependent longevity and thermoresistance. Neuronal HLH-30/TFEB mediated thermoresistance via a W06A11.1-mediated peripheral induction of mitochondria fragmentation in the muscles through a neurotransmission-modulatory mechanism, highlighting a beneficial role of mitochondrial fragmentation in heat stress protection.

## RESULTS

### Neuronal HLH-30/TFEB regulates IIS-dependent longevity

To investigate the activity of HLH-30/TFEB in neurons, we initially directed panneuronal *hlh-30::GFP* reporter expression in wildtype *C. elegans* with the *rab-3* neuronal promoter and the somatic expression-permitting *unc-54* 3’ untranslated region (UTR) (Kimble and Crittenden, 2007). Corroborating previous reports, we observed misregulation of intestinal transgene expression by this promoter and *3’UTR* combination (Wang *et al.*, 2017; Gelino *et al.*, 2016; Silva-García *et al.*, 2019) by visualizing intestinal nuclear enrichment of HLH-30::GFP from *xpo-1* knockdown (Silvestrini *et al.*, 2018), and further discovered that dysregulated expression occurred throughout intestinal tissues and not just posteriorly as previously observed (Figure 1a). We subsequently drove neuronal *hlh-30::GFP* reporter expression with the *rab-3* promoter and its corresponding *rab-3* 3’UTR *(Silva-García et al., 2019)* and confirmed that the *rab-3* 3’UTR was far superior to the widely-used *unc-54* 3’UTR in neuronally-restricting HLH-30::GFP, as reporter expression was only detected in neurons in the absence of intestinal nuclear enrichment from *xpo-*1 knockdown (Figure 1a). We first investigated if neuronal HLH-30/TFEB plays a role in lifespan regulation. In corroboration with previous observations (Lin *et al.*, 2018; Visvikis *et al.*, 2014), *hlh-30(tm1978)* mutants exhibited reduced lifespan in comparison to wildtype animals at 25°C (Figure 1b, Supplemental Figure 1 **and Table 1)**. Additionally, neuronal HLH-30/TFEB rescue in these mutants conferred no significant lifespan improvements in several transgenic lines, apart from one line whereby lifespan was extended which may be attributed to higher expression levels of neuronal HLH-30/TFEB (Figure 1b, Supplemental Figures 1 and 2c, Supplemental Tables 1 **and** 4). Altogether, these observations suggest that re-expressing HLH-30/TFEB in the neurons is not necessarily sufficient to restore normal lifespan.

**Figure 1.**
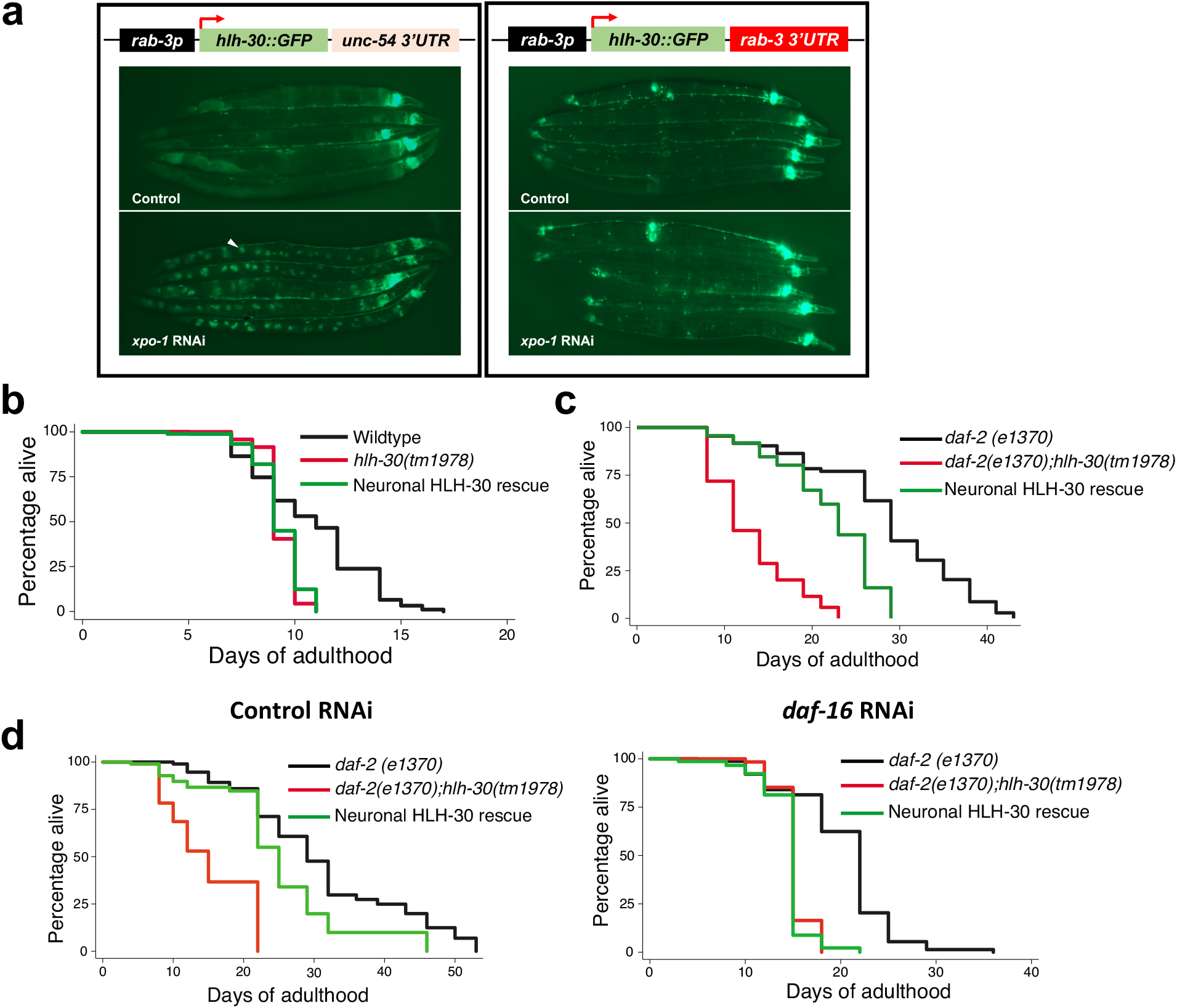
Neuronal HLH-30/TFEB is a regulator of longevity. **(a) (Left panel)** Presence and **(Right panel)** absence of HLH-30::GFP intestinal misexpression (arrowhead, GFP-enriched intestinal nucleus) under combinatorial regulation from the *rab-3* promoter (*rab-3p*) and *unc-54* or *rab-*3’ untranslated region (*3’ UTR*) respectively, as detected with RNAi knockdown of the nuclear exportin gene *xpo-1*, **(b)** Lifespan analyses of wildtype, *hlh-30(tm1978)*, and neuronal HLH-30/TFEB rescued animals fed OP50 at 25°C. **(c and d)** Lifespan analyses of *daf-2(e1370), daf-2(e1370);hlh-30(tm1978)*, and neuronal HLH-30/TFEB rescued animals fed **(c)** OP50 and **(d)** control RNAi (*L4440*) or RNAi against *daf-16* at 25°C. Animals were developed on OP50 at 20°C and shifted to 25°C on **(a to c)** OP50 or **(d)** bacteria expressing either control dsRNA (*L4440*) or dsRNA against *daf-16* from day 1 of adulthood. Data are representatives of 3 independent replicates and comparisons were made by Mantel-Cox log-rank. See Supplemental Figures 1 and 2, and Tables 1 – 3 for additional details on lifespan analyses.

We next investigated the role of neuronal HLH-30 in regulating longevity since systemic HLH-30/TFEB was previously demonstrated to be essential for lifespan extension in several longevity paradigms (Lin *et al.*, 2018; Lapierre *et al.*, 2013). As DAF-16/FOXO was previously reported to co-regulate IIS-dependent longevity with HLH-30/TFEB (Lin *et al.*, 2018), we investigated this in long-lived *daf-2(e1370)* IIS mutants in which longevity is mediated by DAF-16/FOXO activity (Kenyon *et al.*, 1993; Lin *et al.*, 1997). *daf-2(e1370);hlh-30(tm1978)* double mutants exhibited reduced lifespan in comparison to their long-lived *daf-2(e1370)* counterparts, in line with previous studies of *hlh-30* loss of function and silencing in *daf-2* mutants (Figure 1c, Supplemental Figures 2a and b**, Table 2)** (Lapierre *et al.*, 2013; Lin *et al.*, 2018). In contrast to observations in *hlh-30(tm1978)* mutants (Fig. 1b), neuronal HLH-30/TFEB rescue in *daf-2(e1370);hlh-30(tm1978)* animals caused robust and partial rescues of lifespan in several lines (Figure 1c, Supplemental Figures 2a and b**, Table 2)** which was abolished with RNAi-mediated knockdown of *daf-16* (Figure 1d, Supplemental Table 3). As *daf-16* mRNA silencing was peripheral due to the RNAi-refractory nature of neurons in *C. elegans* (Kamath *et al.*, 2001; Timmons, Court and Fire, 2001), this indicates that neuronal HLH-30/TFEB requires peripheral DAF-16/FOXO to regulate IIS-dependent longevity. However, peripheral activation of DAF-16/FOXO from *daf-2* knockdown in wildtype counterparts with normal lifespans did not preserve the lifespan extension mediated by neuronal HLH-30/TFEB rescue in *daf-2(e1370);hlh-30(tm1978)* mutants (Supplemental Figure 2c **and Table 4).** This suggests that neuronal DAF-16/FOXO is a prerequisite for neuronal HLH-30/TFEB for regulating IIS-dependent longevity through peripheral DAF-16/FOXO since *daf-2* silencing is peripheral given the insensitivity of neurons to RNAi, and unlikely to induce neuronal DAF-16/FOXO activity. Taken together, these results demonstrated that neuronal HLH-30/TFEB is not necessarily required for the regulation of normal lifespan but is essential for IIS-dependent longevity.

### Neuronal HLH-30/TFEB mediates heat stress resistance independently of DAF-16/FOXO

The relationship between thermoresistance and lifespan extension was demonstrated by improved thermotolerance from pro-longevity mutations and lifespan-extending effects of transient heat stress exposure in *C. elegans* (Kumsta *et al.*, 2017; Lithgow *et al.*, 1995; Butov *et al.*, 2001; Michalski *et al.*, 2001). Since HLH-30/TFEB is systemically required for thermotolerance (Lin *et al.*, 2018) and long-lived *daf-2(e1370)* mutants exhibited improved thermoresistance than wildtype animals (Lithgow *et al.*, 1995), our observations of neuronal HLH-30/TFEB-regulated IIS-dependent longevity prompted us to investigate whether it also mediates thermoresistance. In animals expressing the HLH-30::GFP reporter either ubiquitously or neuronally, exposure to acute heat stress at 37°C induced HLH-30/TFEB nuclear enrichment (Figure 2a), in line with previous observations (Lin *et al.*, 2018). Furthermore, nuclear enrichment was augmented with prolonged heat stress, suggesting an increased requirement of the transcriptional activity of HLH-30/TFEB to counteract heat stress. Notably, *hlh-30(tm1978)* mutants exhibited compromised survival in comparison to wildtype animals with prolonged heat stress, which was mitigated by the neuronal rescue of HLH-30/TFEB (Figure 2b, Supplemental Figure 3a **and** b). Surprisingly, we found that neuronal HLH-30/TFEB rescue failed to improve the reduced survival of *daf-2(e1370);hlh-30(tm1978)* double mutants to heat stress (Figure 2c, Supplemental Figures 3c **and** d). In sum, our observations demonstrate that neuronal HLH-30/TFEB is important for heat stress resistance in wildtype but not in long-lived *daf-2(e1370)* animals.

**Figure 2.**
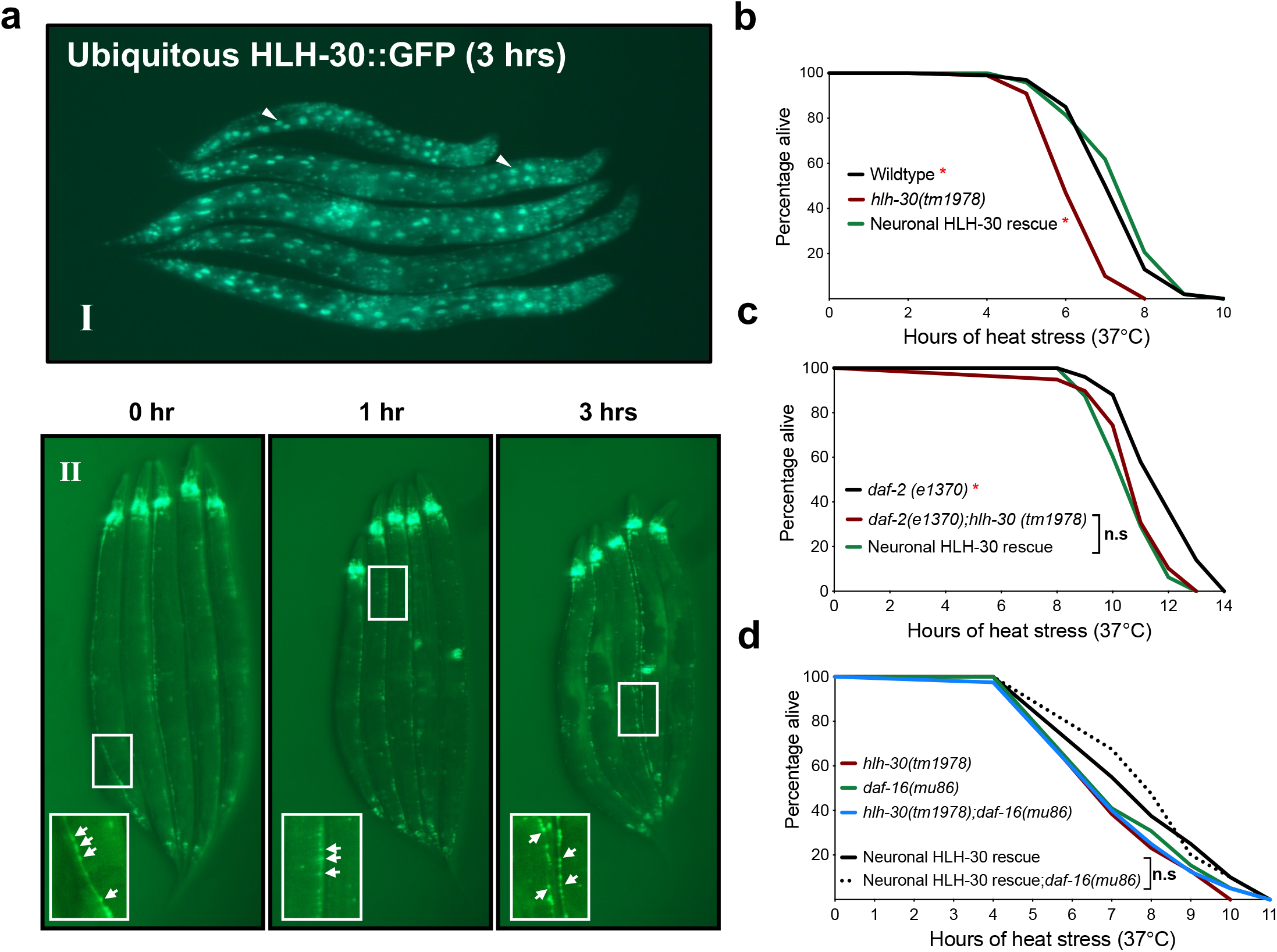
Neuronal HLH-30/TFEB mediates resistance to heat stress. **(a)** Visualization of **(I)** ubiquitous (arrowheads, intestinal cells) and **(II)** panneuronal (arrows, enlarged insets of ventral nerve cords) HLH-30::GFP nuclear enrichment following 37°C heat stress for up to 3 hrs. **(b and c)** Survival analyses of neuronal HLH-30/TFEB rescued animals in comparison with their **(b)** wildtype, *hlh-30(tm1978),* **(c)** *daf-2(e1370),* and *daf-2(e1370);hlh-30(tm1978)* counterparts at 37°C heat stress. **(d)** Survival analyses of neuronal HLH-30/TFEB rescued *hlh-30(tm1978)* animals in in the presence and absence of *daf-16(mu86)* loss of function at 37°C heat stress. Animals were developed at 20°C and shifted to 37°C to induce heat stress at day 1 of adulthood. Data are representatives of 2 - 4 independent replicates and comparisons were made by Mantel-Cox log-rank (*n* = 78 - 200/strain; n.s, *p*≥0.05; *, *p*<0.05; in comparison to *hlh-30(tm1978)* or *daf-2(e1370);hlh-30(tm1978)*). See Supplemental Figure 3 for heat stress survival analyses on additional neuronal HLH-30/TFEB rescued lines.

Interestingly, our findings indicate that there are differential requirements for neuronal HLH-30/TFEB in regulating longevity and thermoresistance in wildtype and long-lived genetic backgrounds. Moreover, the lack of survival improvements with neuronal HLH-30/TFEB rescue in *daf-2(e1370);hlh-30(tm1978)* double mutants despite DAF-16/FOXO-dependent lifespan extension suggests that neuronal HLH-30/TFEB does not regulate thermoresistance through DAF-16/FOXO, in line with previous findings that these transcription factors mediate heat stress protection through separate pathways (Lin *et al.*, 2018). Indeed, loss of function *daf-16(mu86)* did not dampen the survival of neuronal HLH-30/TFEB rescued animals to heat stress in non-*daf-2(e1370)* animals (Figure 2d). Taken together, these findings demonstrated that neuronal HLH-30/TFEB regulates longevity and thermoresistance dependently and independently of DAF-16/FOXO respectively.

### Neuronal HLH-30/TFEB mediates thermoresistance via the protein W06A11.1

To gain insights into mechanisms mobilized by neuronal HLH-30/TFEB to mediate heat stress resistance, we compared RNAseq analyses of wildtype, *hlh-30(tm1978)*, and neuronal HLH-30/TFEB rescued animals from control and heat stressed conditions. We identified heat stress-induced differentially expressed genes (DEGs) which were up-and downregulated for each genotype and focused on significant expression changes shared by wildtype and neuronal HLH-30/TFEB rescued animals in comparison to *hlh-30(tm1978)* mutants. Heat stress-induced DEGs were overlapped between the genotypes to extract those unique to wildtype and neuronal HLH-30/TFEB rescued animals, or which exhibited a greater expression change in those genotypes in comparison to *hlh-30(tm1978)* mutants (Figure 3a **and** Supplemental Figure 4). From these analyses, we applied thresholds for adjusted *p*-values of significance and Log_2_ fold changes and derived a total of 25 upregulated and downregulated DEGs (Figure 3a **and** Supplemental Figure 4).

**Figure 3.**
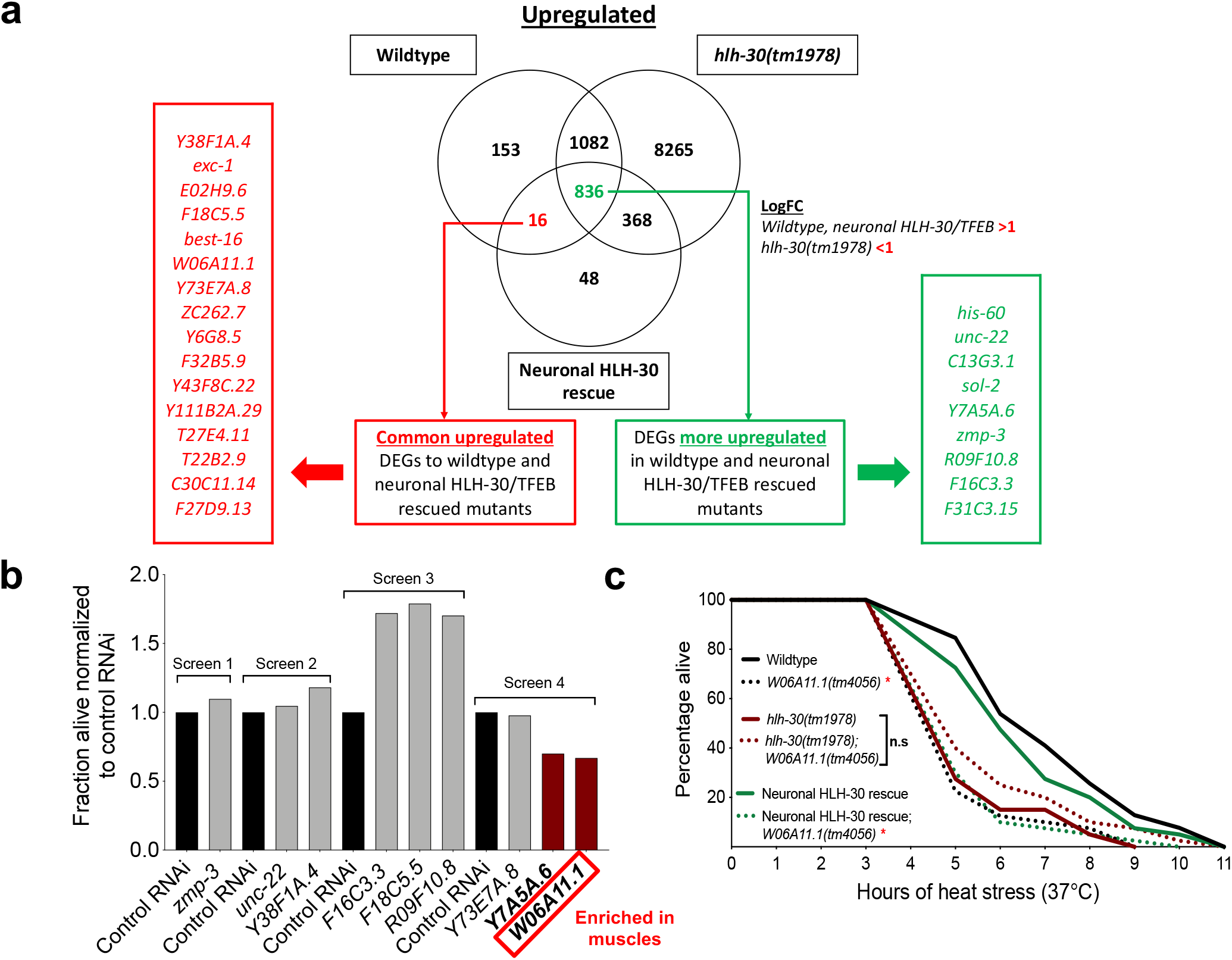
*W06A11.1* is a novel and bona fide mediator of thermoresistance. **(a)** Pairwise comparisons of RNAseq profiles of day 1 animals from 20°C (control conditions) and 3 hrs heat stress at 37°C identified genotype-specific differentially expressed genes (DEGs) upregulated with heat stress, which were overlapped between genotypes to extract significant hits (adjusted *p*<0.05) common to or more upregulated (Log_2_ fold change (LogFC) thresholds applied as indicated) in wildtype and neuronal HLH-30/TFEB rescued animals than *hlh-30(tm1978)* mutants. Animals from 4 independent replicates were developed at 20°C to day 1 of adulthood and harvested for RNA after 3 hrs of further growth at 20°C or 37°C. **(b)** Survival analyses of neuronal HLH-30/TFEB rescued animals after 7 hrs heat stress at 37°C following RNAi knockdown of upregulated hits. **(c)** Survival analyses of wildtype, *hlh-30(tm1978)*, and neuronal HLH-30/TFEB rescued animals in the absence and presence of *W06A11.1(tm4056)* loss of function at 37°C heat stress. **(b and c)** Animals were developed at 20°C to day 1 of adulthood and exposed to 37°C heat stress for **(b)** 7 hrs after feeding with RNAi bacteria for 48 hrs or **(b)** until death. Data are representatives of **(c)** single experiments (*n* = 48 - 53/RNAi; fraction alive normalized to control RNAi per screen) and **(c)** 2 independent replicates (comparisons by Mantel-Cox log-rank; *n* = 79 - 80/strain; n.s, *p*≥0.05; *, *p*<0.05; *W06A11.1(tm4056)* compared to control per genotype).

In order to uncover new modulators of heat stress resistance via neuronal HLH-30/TFEB, gene knockdowns were performed on upregulated DEGs with available RNAi clones in initial screens with neuronal HLH-30/TFEB rescued animals to functionally test their requirement for heat stress resistance (Figure 3b). Two hits which dampened survival to heat stress with knockdown were identified, *Y7A5A.6* and *W06A11.1*, which have uncharacterized functions but were previously found to be enriched in subsets of neurons (Lockhead *et al.*, 2016). We followed up on W06A11.1 since its expression is also enriched in the muscle (Fox *et al.*, 2007), a tissue that can modulate lifespan and proteostasis in response to neuronal signaling (Garcia *et al.*, 2007; Tatum *et al.*, 2015; Prahlad and Morimoto, 2011; Silva, Amaral and Morimoto, 2013; O’Brien *et al.*, 2018; van Oosten-Hawle, Porter and Morimoto, 2013; Zhang *et al.*, 2019; Burkewitz *et al.*, 2015). Of note, our findings of *hlh-30*-dependent upregulation of *W06A11.1* with heat stress corroborated previous transcriptomics comparisons between heat stressed wildtype and *hlh-30(tm1978)* mutants (Lin *et al.*, 2018). We found that both loss of function *W06A11.1(tm4056)* and RNAi-mediated peripheral knockdown of *W06A11.1* compromised the survival of wildtype and neuronal HLH-30/TFEB rescued animals but not of *hlh-30(tm1978)* mutants in heat stress (Figures 3c **and** Supplemental Figure 5a). These findings confirmed that W06A11.1 is a bona fide mediator of thermoresistance and support the potential occurrence of inter-tissue signaling events between neuronal HLH-30/TFEB and distal W06A11.1.

### Neuronal HLH-30/TFEB induces mitochondrial fragmentation in muscles via W06A11.1 to improve thermoresistance

Heat stress was previously found to cause increased mitochondrial fragmentation in muscles (Momma *et al.*, 2017; Machiela *et al.*, 2020; Chen *et al.*, 2021), and disrupting mitochondrial fission and fusion were detrimental and beneficial to heat stress resistance respectively, suggesting that mitochondrial fragmentation is mechanistically important for thermoresistance (Machiela *et al.*, 2020). Given the muscle-enriched expression of *W06A11.1* (Fox *et al.*, 2007), and that neuronal signaling can induce distal mitochondrial fragmentation in muscles (Burkewitz *et al.*, 2015; Zhang *et al.*, 2019), we wondered if neuronal HLH-30/TFEB mediates thermoresistance through W06A11.1-dependent muscle mitochondrial fragmentation. To investigate this, we first confirmed the functional importance of mitochondrial fragmentation in heat stress resistance as knocking down the mitochondrial fission and fusion genes, *drp-1* and *eat-3,* respectively dampened and improved survival of wildtype animals to heat stress (Figure 4a and b). We then utilized a body wall muscle-targeted, GFP-tagged mitochondrial reporter (Mito::GFP) (Sarasija and Norman, 2015) to compare mitochondrial morphology between the three genotypes. Corroborating previous observations (Momma *et al.*, 2017; Machiela *et al.*, 2020; Chen *et al.*, 2021), muscle mitochondria were reticular at 20°C (Figure 4cI) but exhibited extensive fragmentation following heat stress (Figure 4cII). As mitochondrial fragmentation often preludes cell death (Suen, Norris and Youle, 2008), we investigated if heat stress had additionally induced muscle degeneration. We observed that the continual striated morphology of myosin heavy chain filaments was not compromised by heat stress, using a MYO-3::GFP-expressing strain (Campagnola *et al.*, 2002) (Figure 4cIV).

**Figure 4.**
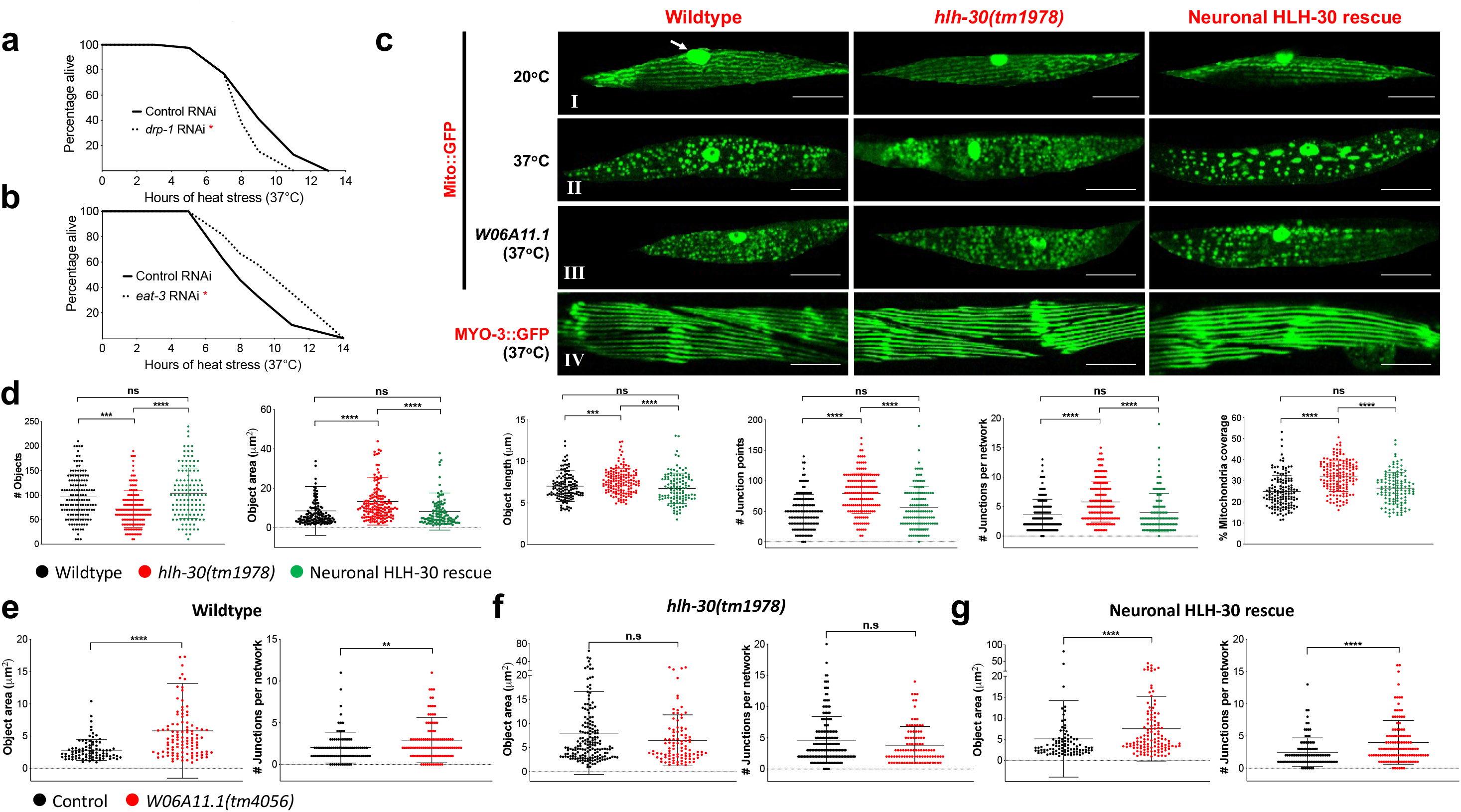
Neuronal HLH-30/TFEB mediates thermoresistance through non-cell autonomous induction of *W06A11.1*-dependent mitochondrial fragmentation in muscles. **(a and b)** Survival analyses of wildtype animals fed control RNAi (*L4440*) or RNAi against **(a)** *drp-1* and **(b)** *eat-3* at 37°C heat stress. Animals developed at 20°C to the L4 larval stage on OP50 were transferred onto bacteria expressing either control dsRNA (*L4440*) or dsRNA against *drp-1* or *eat-3*, grown at 25°C for 48 hrs, and exposed to 37°C heat stress. Data are representatives of 2 independent replicates (*n* = 89 - 180/RNAi) and comparisons were made by Mantel-Cox log-rank (*, *p*<0.05; in comparison to control RNAi). **(cI to III)** Representative images of muscle mitochondrial morphology with the body wall muscle mitochondrial reporter (Mito::GFP) in wildtype, *hlh-30(tm1978)*, and neuronal HLH-30/TFEB rescued animals at **(I)** 20°C (arrow, GFP in muscle nuclei) and after 37°C heat stress for 3 hrs in the **(II)** absence and **(III)** presence of *W06A11.1(tm4056)* loss of function. **(cIV)** Representative images of muscle morphologies with the body wall muscle myosin reporter (MYO-3::GFP) after 37°C heat stress for 3 hrs. **(d)** Analysis of mitochondrial connectivity in wildtype, *hlh-30(tm1978*, and neuronal HLH-30/TFEB rescued animals after 37°C heat stress for 3 hrs. **(e to g)** Analysis of mitochondrial connectivity in the absence and presence of *W06A11.1(tm4056)* loss of function after 37°C heat stress for 3 hrs in **(e)** wildtype, (f) *hlh-30(tm1978)* loss of function mutants, and **(g)** neuronal HLH-30/TFEB rescued animals (see Supplemental Figure 5c to d for additional mitochondrial quantification). **(c to e)** Animals were developed at 20°C to day 1 of adulthood and exposed to 37°C heat stress for 3 hrs. Data are representatives of **(d)** 4 independent replicates (per strain; *n* = 40, number of ROIs = 117 – 150), and **(e to g)** 3 independent replicates (per strain; *n =* 30, number of ROIs = 88 – 171). Comparisons were made by **(d)** Kruskal-Wallis or **(e to g)** Mann-Whitney or student’s unpaired *t-*test accordingly to the normality of data distribution for each analyzed mitochondrial feature (presented as mean ± S.D; n.s, *p*≥0.05; *, *p*<0.05; **, *p*<0.01; ***, *p*<0.001; ****, *p*<0.0001). Scale bars = 20 μM.

In order to compare the extents of heat stress-induced mitochondrial fragmentation between the genotypes, we further quantified various parameters of mitochondrial morphology using MitoMAPR (Zhang *et al.*, 2019). Strikingly, we found that heat stressed *hlh-30(tm1978)* mitochondria had lower counts but higher areas and length, suggesting reduced breakdown of mitochondrial tubularity (Figure 4d). Furthermore, increased junction points, junctions per mitochondrial network, and mitochondrial coverages, suggest higher mitochondrial network connectivity in these mutants (Figure 4d). Altogether, this indicates that heat stress-induced mitochondrial fragmentation in the muscles of *hlh-30(tm1978)* mutants were reduced in comparison to wildtype and neuronal HLH-30/TFEB rescued animals. In addition to our observations that mitochondrial fission was required for survival to heat stress (Figure 4a and b), this supports that neuronal HLH-30/TFEB mediates thermoresistance by inducing mitochondrial fragmentation in distal muscles.

In order to assess the contribution of W06A11.1 in heat stress-induced mitochondrial fragmentation, we tested *W06A11.1(tm4056)* mutants. We found that the loss of W06A11.1 function repressed mitochondrial fragmentation during heat stress in wildtype and neuronal HLH-30/TFEB rescued animals whilst having no effect in *hlh-30(tm1978)* mutants (Figures 4cIII, e **to** g, Supplemental Figures 5b **to** d). Taken together, our findings demonstrate that neuronal HLH-30/TFEB non-cell autonomously regulates heat stress resistance by inducing muscle mitochondrial fragmentation via W06A11.1 function.

### Neuronal HLH-30/TFEB induces distal muscle mitochondrial fragmentation by modulating neurotransmission to mediate thermoresistance

Since neuronal HLH-30/TFEB mediates the induction of muscle mitochondrial morphology changes during heat stress, we hypothesized that neurotransmission or neurosecretion signaling events were regulated by HLH-30/TFEB. To test this, we leveraged the *unc-13(e1091)* and *unc-31(e928)* loss of function mutations which result in defective synaptic vesicle and dense core vesicle (DCV) release respectively (Richmond, Davis and Jorgensen, 1999; Speese *et al.*, 2007). We found that defective DCV release universally improved the thermoresistance of all three genotypes (Supplemental Figure 6), demonstrating that neurosecretory events are HLH-30/TFEB-independent and detrimental for heat stress resistance, in line with a previous study showing a deleterious effect of neurosecretory signaling on proteostasis in the muscle (Prahlad and Morimoto, 2011). Additionally, mitochondrial fragmentation was not affected by defective DCV release, indicating that antagonism of heat stress resistance by neurosecretory signaling occurs via a separate mechanism (Supplemental Figure 7).

In contrast, defective synaptic vesicle release markedly improved the thermoresistance of *hlh-30(tm1978)* mutants (Figure 5a), which corresponded with enhanced heat stress-induced mitochondrial fragmentation in muscles (Figure 5b and d, Supplemental Figure 8b). Notably, loss of neurotransmission impacted mitochondrial fragmentation in wildtype and neuronal HLH-30/TFEB rescued animals although thermoresistance was not significantly affected, suggesting mechanistic overlap between the loss of *unc-13* and neuronal HLH-30/TFEB (Figure 5a to e, Supplemental Figure 8a **and** c). Overall, these findings suggest that neurotransmission signaling impairs thermoresistance. Additionally, our findings demonstrate that neurotransmission signaling regulates distal induction of mitochondrial fragmentation in muscles. Taken together, this suggests that neuronal HLH-30/TFEB mediates thermoresistance through neurotransmission to induce distal mitochondrial fragmentation in muscles.

**Figure 5.**
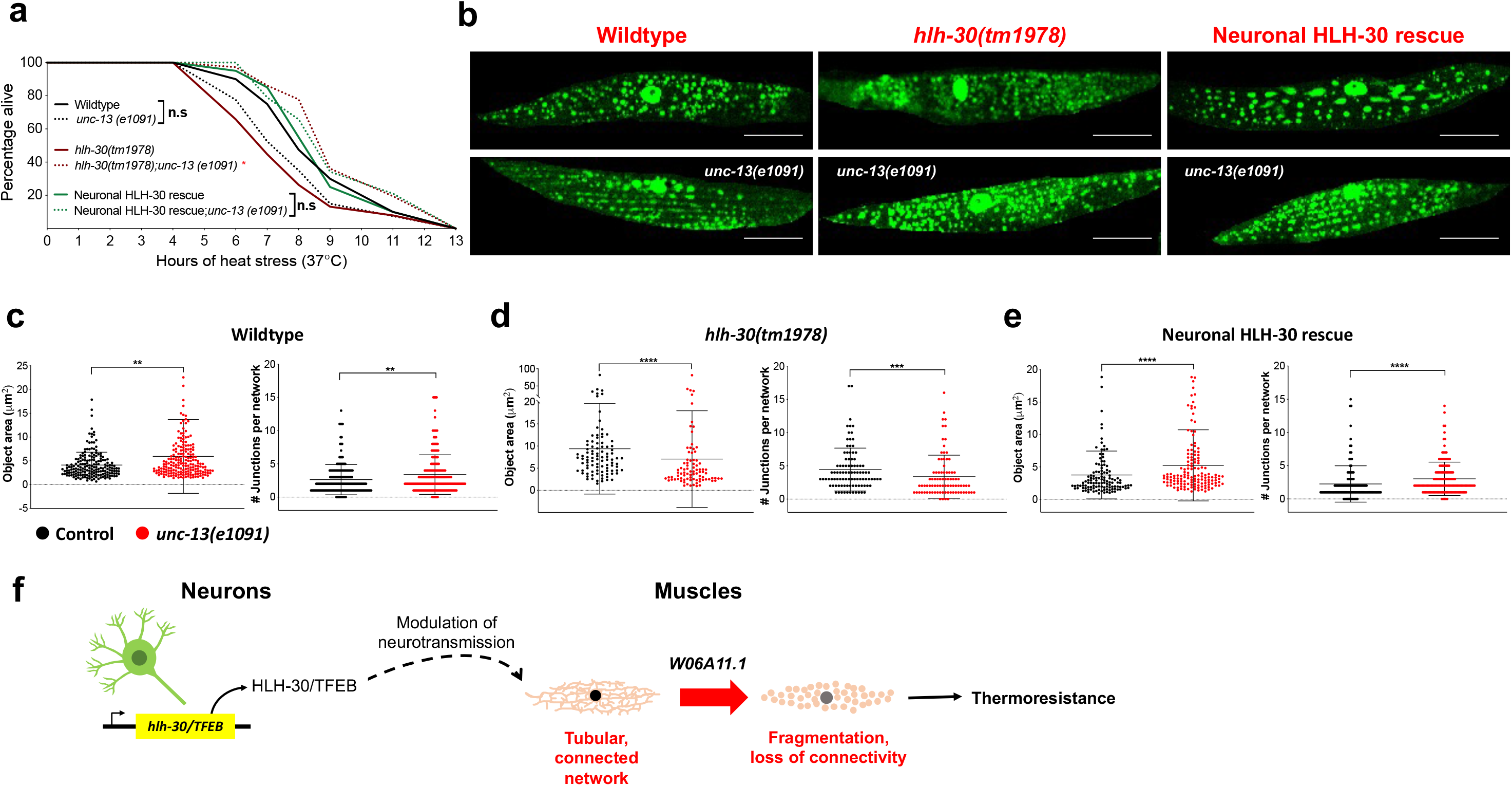
Neuronal HLH-30 induces peripheral muscle mitochondria fragmentation by modulating neurotransmission during heat stress. **(a)** Survival analyses of wildtype, *hlh-30(tm1978)*, and neuronal HLH-30/TFEB rescued animals in the absence and presence of *unc-13(e1091)* loss of function at 37°C heat stress. Animals were developed at 20°C to day 1 of adulthood and exposed to 37°C heat stress. Data is representative of 3 - 4 independent replicates (*n* = 115 - 170/strain) and comparisons were made by Mantel-Cox log-rank (n.s, *p*≥0.05; *,*p*<0.05; comparisons of *unc-13(e1091)* to control per genotype). **(b)** Representative images of muscle mitochondrial morphology with the body wall muscle mitochondrial reporter (Mito::GFP) in wildtype, *hlh-30(tm1978)*, and neuronal HLH-30/TFEB rescued animals in the **(top panel)** absence and **(bottom panel)** presence of *unc-13(e1091)* loss of function after 37°C heat stress for 3 hrs. **(c to e)** Analysis of mitochondrial connectivity in the absence and presence of *unc-13(e1091)* loss of function in **(c)** wildtype, **(d)** *hlh-30(tm1978)*, and **(e)** neuronal HLH-30/TFEB rescued animals after 37°C heat stress for 3 hrs (see Supplemental Figure 8 for additional mitochondrial quantification). **(b to e)** Animals were developed at 20°C to day 1 of adulthood and exposed to 37°C heat stress for 3 hrs. Data are representatives of 3 - 5 independent replicates (per strain; *n* = 30 - 50; number of ROIs = 87 – 196) and comparisons were made by either Mann-Whitney or student’s unpaired *t-*test accordingly to the normality of data distribution for each analyzed mitochondrial feature (presented as mean ± S.D; n.s, *p*≥0.05; *, *p*<0.05; **, *p*<0.01; ***, *p*<0.001; ****, *p*<0.0001). Scale bars = 20 μM. **(f)** Model depicting non-cell autonomous regulation of thermoresistance by neuronal HLH-30/TFEB.

## DISCUSSION

A mechanistic understanding of lifespan and stress resistance regulation by TFEB, as well as its cell-and non-cell autonomous-associated mechanisms, is crucial to properly leverage TFEB activity to promote healthy aging. Our study provides pioneering evidence that HLH-30/TFEB in neurons regulates heat stress resistance and IIS-related longevity by distinct mechanisms. Importantly, we report that neuronal HLH-30/TFEB non-cell autonomously regulates heat stress resistance by modulating neurotransmission, in turn stimulating mitochondrial fragmentation distally in the muscles of *C. elegans* through the protein, W06A11.1 (Figure 5f).

Previous studies have demonstrated that HLH-30/TFEB is systemically required for maintaining normal lifespan, longevity, and heat stress resistance in *C. elegans* (Visvikis *et al.*, 2014; Lapierre *et al.*, 2013; Lin *et al.*, 2018). As neurons are important for integrating signaling events to regulate these events (Miller *et al.*, 2020), we hypothesized that HLH-30/TFEB activity in neurons may be similarly important for upkeeping these processes. Notably, we observed that neuronal HLH-30/TFEB is essential for IIS-dependent longevity but redundant for thermoresistance in long-lived *daf-2* mutants. Conversely, it is not necessarily required for normal lifespan but important for thermoresistance in non-long-lived animals. These diametrically opposite requirements of neuronal HLH-30/TFEB for lifespan and heat stress resistance in longevity-promoting and non-promoting genetic backgrounds highlight a potential divergence of these regulatory mechanisms. Indeed, they were uncoupled with DAF-16/FOXO, which was required by neuronal HLH-30/TFEB for the promotion of longevity but not for thermoresistance. This corroborated separate findings of context-dependent synergy between HLH-30/TFEB and DAF-16/FOXO for the regulation of longevity but not heat stress resistance (Lin *et al.*, 2018). Notably, previous work has shown that the neuronal activity of another transcription factor, HSF-1, can also employ separate mechanisms to regulate longevity and heat stress resistance (Douglas *et al.*, 2015). Hence, our findings provide further compelling evidence that transcription factors can employ context-specific mechanisms to regulate systemic processes from neurons. Additionally, our observations support that the association of enhanced stress resistance with longevity is largely contextual (Dues *et al.*, 2019; Dues *et al.*, 2017). Taken together, this places emphasis on the importance of dissecting the diverse tissue-specific mechanisms of longevity-and stress-modulating proteins such as HLH-30/TFEB to better target age-associated decline.

Transcriptional responses to heat stress, such as HSF1-dependent upregulation of the heat shock response, are important in enabling organisms to counteract heat-associated proteostatic insults (Morimoto, 2011; Rodriguez *et al.*, 2013). Herein, we identified heat-induced, neuronal HLH-30/TFEB-dependent transcriptional changes which were similarly preserved in the presence of ubiquitous HLH-30/TFEB, suggesting that neuronal HLH-30/TFEB markedly affects peripheral gene expression and/or processes to bring about detectable transcriptional changes at the systemic level. We surprisingly found a downregulation of *hlh-30/TFEB* transcripts with heat stress which was not attributed to reduced nuclear entry (Figure 2a, Supplemental Figure 4), suggesting possible negative autoregulation which is reported to be important for speeding transcriptional responses (Rosenfeld, Elowitz and Alon, 2002). Most importantly, we found that neuronal HLH-30/TFEB mediates thermoresistance dependently on the gene *W06A11.1*, by communicating non-cell autonomously through neurotransmission inhibition to induce peripheral W06A11.1-dependent mitochondrial fragmentation in muscles. These observations raise the question of how W06A11.1 mediates mitochondrial fragmentation. The mechanistic functions of W06A11.1 are uncharacterized, but it may be directly involved in mitochondrial fragmentation akin to the mitochondrial fission factor DRP-1, or indirectly, such as by post-translationally activating DRP-1 or acting as mitochondrial membrane-bound factors which recruit DRP-1 to scission sites (Sharma *et al.*, 2019). In the same vein, the mechanisms mobilized by neuronal HLH-30/TFEB for modulating neurotransmission to influence peripheral W06A11.1-dependent mitochondrial fragmentation in muscles are unclear, but observations herein suggest that hyperactivated neuronal signaling to the periphery impairs heat stress resistance, resulting in heat-induced proteostatic challenges. Notably, imbalanced neuronal hyperstimulation due to excessive cholinergic excitation and loss of gamma-aminobutyric acid inhibition at neuromuscular junctions was shown to be detrimental for proteostasis in muscles (Garcia *et al.*, 2007), lending support to this hypothesis. Going forward, it is important to elucidate the neuromodulatory mechanisms which are dependent on neuronal HLH-30/TFEB at neuromuscular junctions to further understand the mechanistic processes leading to peripheral mitochondrial fragmentation in the muscles.

An unexpected finding of our study is the protective role of mitochondrial fragmentation against heat stress. In contrast, mitochondrial fragmentation was previously shown to be detrimental for longevity (Burkewitz *et al.*, 2015; Zhang *et al.*, 2019; Gerisch *et al.*, 2020; Zhou *et al.*, 2019). Of note, previous observations associating adult reproductive diapause-induced longevity with HLH-30/TFEB-dependent inhibition of muscle mitochondrial fragmentation (Gerisch *et al.*, 2020) demonstrated a reverse role of HLH-30/TFEB in promoting mitochondrial fusion rather than fragmentation to mediate longevity, which suggests context-specific mitochondrial dynamics regulation by HLH-30/TFEB. As mitochondria propagate diverse types of signals which are regulated by their dynamics (Chandel, 2015; Labbé, Murley and Nunnari, 2014), heat stress protection may potentially be mediated by any of these downstream mitochondrial related signaling events. Notably, mitochondrial fragmentation induced by cell injury was shown to be important for mediating localized plasma membrane repair through mitochondrial calcium-dependent redox signaling, exemplifying a protective role of mitochondrial fragmentation for stress mitigation (Horn *et al.*, 2020). Altogether, our work revealed a novel non-cell autonomous mechanism of thermoresistance originating in neurons, and puts neuronal HLH-30/TFEB as a target of interest to enable systemic improvement in proteostasis to promote healthy aging.

## METHODS

### C. elegans strains

All strains were cultured on *Escherichia coli* OP50 on nematode growth media (NGM) agar at 20°C as previously described (Stiernagle, 2006; Brenner, 1974) (additional strain details provided in Supplemental Table 5). Synchronous populations were obtained either by sodium hypochlorite bleaching or egg picking.

### Generation of neuronal HLH-30::GFP DNA construct

The DNA construct pLP27 (*rab-3p::hlh-30::GFP::3XFLAG::rab-3)* was designed to neuronally-restrict *hlh-30* transgene expression with the promoter and *3’UTR* regulatory elements of the neuronal *rab-3* gene. pLP27 was generated by replacing the *unc-54 3’UTR* in pLP21 (*rab-3p::hlh-30::GFP::3XFLAG::unc-54*). The following describes the sequential generation of pLP21, followed by pLP27. The 3XFLAG-tag was cloned from pLP15 (Addgene plasmid #55180) with 5’ KpnI and 3’ EcoRI sites (primers LC127 and 128) into the corresponding restriction digestion sites of pLP9 (Addgene plasmid #1497, Fire Lab vector kit construct pPD95.81) to generate pLP16 (pPD95.81_*3XFLAG*). The *hlh-30* coding sequence was then cloned from *C. elegans* cDNA with 5’ KpnI and 3’ NaeI sites (primers LC154 and 155) and inserted upstream of the 3XFLAG-tag into the corresponding restriction digestion sites of pLP16. The construct was cloned as linearized vectors with vector-specific primers LC184 and 185 for insertion of the *GFP* sequence (cloned from pLP19 (*hlh-17p::GFP*) with insert-specific primers LC186 and 187) between *hlh-30* and *3XFLAG*-tag segments with HiFi cloning (NEBuilder® HiFi DNA Assembly Cloning Kit, New England BioLabs Inc., Ipswich, MA). Lastly, the *rab-3* gene promoter was cloned from pLP17 (*rab-3p::sid-1*) with 5’ SphI and 3’ KpnI sites (primers LC143 and 188) into the corresponding restriction digestion sites of the preceding construct to generate pLP21. To derive pLP27 from pLP21, the *rab-3* gene *3’ UTR* was amplified from *C. elegans* genomic DNA (insert-specific primers LC227 and 228) and inserted into pLP21 in place of *unc-53 3’UTR* (vector-specific primers LC225 and 226) with HiFi cloning. Primers used are listed in Supplemental Table 6. pLp17 (plasmid #462) and pLP19 (plasmid #836) were kindly provided by Dr. Andrew Dillin (UC Berkeley, CA).

### Construction of neuronal HLH-30::GFP transgenic strains

Transgenic strains were generated by microinjecting the germlines of day 1 wildtype N2 or *hlh-30(tm1978)* adults with 10 ng/uL pLp11 (*hlh-30p::hlh-30::GFP::unc-54*) (Lapierre *et al.*, 2013) or 25 ng/uL pLP27 (*rab-3p::hlh-30::GFP::3XFLAG::rab-3 3’UTR*) with 85 ng/uL of pLP24 (*unc-122p::RFP)* as a selection marker; pLP24 was obtained from Addgene (plasmid #8938) (Miyabayashi et al., 1999). Where indicated in Supplemental Table 5, transgenic progenies were integrated for extrachromosomal arrays by UV irradiation, and further outcrossed to wildtype animals for ten times. The expression and nuclear localization of HLH-30::GFP were visualized by imaging animals immobilized onto NGM agar with 0.1 % sodium azide with a Zeiss Discovery V20 fluorescence microscope (Zeiss, White Plains, NY). To prevent nuclear localization caused by stress from immobilization, imaging was performed within 5 mins of mounting (Lapierre *et al.*, 2013).

### Genotyping

Individual animals were lysed in 10X Standard Taq buffer/proteinase K, and lysates were used as gDNA templates for PCR amplification with OneTaq® Quick-Load® 2X Master Mix with Standard Buffer (all materials from New England BioLabs, Ipswich, MA). Genotyping was performed with primers listed in Supplemental Table 6 with the following thermocycling conditions: Initial denaturation at 95°C (5 mins), 40 cycles of denaturation (95°C, 45s)/annealing (60°C, 45s)/extension (72°C, 1 min/kb product), and final extension at 72°C (10 mins).

### Lifespan analysis

Synchronous populations were developed at 20°C on OP50 until day 1 of adulthood, followed by growth at 25°C on plates containing either OP50 or RNAi clones from the Ahringer library (Kamath *et al.*, 2003). Animals were assessed for their survival every 2-3 days as previously described (Hamilton *et al.*, 2005).

### Analysis of survival to heat stress

Synchronized populations were developed at 20°C were shifted to 37°C for heat stress induction at day 1 of adulthood. For RNAi-mediated knockdown of genes, animals were transferred onto RNAi clones from the Ahringer library (Kamath *et al.*, 2003) from either the L4 larval stage or day 1 of adulthood and maintained at 20°C for 48 hrs, followed by subsequent exposure to 37°C heat stress. Starting from the 3^rd^ or 4^th^ hour of heat stress, the survival of animals was scored every 1 – 3 hrs until all animals were dead.

### RNA sample preparation and RNAseq

Approximately 10,000 to 12,000 synchronized animals developed at 20°C per independent replicate for a total of 4 replicates were washed and either kept at control conditions (20°C) or shifted to 37°C for a non-lethal duration of 3 hrs on day 1 of adulthood. Animals were frozen down at −80°C and subsequently extracted for RNA as previously described (Mills *et al.*, 2019), followed by DNase treatment with the Qiagen RNase-Free DNase Set and purification by the Qiagen RNeasy Mini Kit (Qiagen, Germantown, MD). RNA samples were confirmed for their quality with the Agilent 2100 Bioanalyzer (Agilent Technologies, Santa Clara, CA), and sequenced by GENEWIZ (Azenta Life Sciences, South Plainfield, NJ). Briefly, RNAseq libraries were prepared with the NEBNext Ultra II RNA Library Prep Kit for Illumina (New England BioLabs Inc., Ipswich, MA) and sequenced with the Illumina HiSeq instrument (Illumina Inc., San Diego, CA) with a 2×150bp Paired End configuration. Generated raw sequence files were converted into fastq data, followed by de-multiplexing with the bcl2fastq 2.17 software from Illumina.

### RNAseq data analysis

RNAseq outputs were further subjected to differential expression analysis by GENEWIZ. Briefly, sequence reads trimmed with Trimmomatic v.0.36 (Bolger, Lohse and Usadel, 2014) were mapped to the *C. elegans* ENSEMBL reference genome with STAR aligner v.2.5.2b (Dobin *et al.*, 2013), followed by the quantification of gene counts from the Subread package v.1.5.2. Pairwise comparisons of gene expression between control and heat stressed groups per genotype were subsequently performed to identify heat stress-induced differential gene expressions. Comparisons were carried out with DESeq2 (Love, Huber and Anders, 2014) on raw gene count tables filtered to remove low and non-expressing genes across all samples (average of <10 counts per gene across all samples). Differential expression was performed using the Wald test to calculate Log_2_ fold changes and adjusted *p*-values were generated using Benjamini & Hochberg with an alpha = 0.05. Differential expressions with adjusted *p*-values <0.05 were considered significant.

Heat stress-induced up-and downregulated DEGs for each genotype were further overlapped with Venny (publicly available at http://bioinfogp.cnb.csic.es/tools/venny/index.html) in this study to identify significant DEGs (adjusted *p*<0.05) specific to or with greater extents of change in wildtype and neuronal HLH-30/TFEB animals. Overlapping analysis was performed with significant DEGs from wildtype and neuronal HLH-30/TFEB animals and both significant and non-significant DEGs (adjusted *p*≥0.05) from *hlh-30(tm1978)* mutants. To identify DEGs with greater expression changes in both directionalities in wildtype and neuronal HLH-30/TFEB animals, the following Log_2_FC thresholds were additionally applied: Upregulated (wildtype and neuronal HLH-30/TFEB (Log_2_FC >1), *hlh-30(tm1978)* (Log_2_FC<1)), downregulated (wildtype and neuronal HLH-30/TFEB (Log_2_FC <-1), *hlh-30(tm1978)* (Log_2_FC >-1)).

### Confocal imaging

Day 1 animals expressing GFP-labeled body wall muscles (MYO-3::GFP) or muscle mitochondria (Mito::GFP) at control (20°C) or heat stressed (3hrs at 37°C) were immobilized with 0.1% sodium azide onto 3% agarose pads. 2D images were acquired with Olympus FV3000 confocal laser scanning microscope (Olympus Scientific Solutions Americas Corp., Waltham, MA).

### Analysis of mitochondria morphology

Regions of interest (ROIs, 45 x 45 pixels) from Mito::GFP confocal images were analyzed and quantified for various mitochondrial parameters of network connectivity with the MitoMAPR macro in FIJI (Schindelin *et al.*, 2012) as previously described (Zhang *et al.*, 2019). Quantified outputs provide overall insights into mitochondrial integrity and connectivity and was used in this study to compare extent of fragmentation. Analyzed features are described in more detail previously (Zhang *et al.*, 2019).

### Statistics

Statistical comparisons of survival for lifespan and heat stress were performed with the Mantel-Cox log-rank test with STATA (StataCorp, College Station, TX). Mitochondrial parameters quantified from MitoMAPR analyses were statistically compared with Kruskal-Wallis multiple comparisons test, Mann-Whitney test, or student’s unpaired *t-*test after normality of data distribution were determined with GraphPad Prism 9 (GraphPad Software, San Diego, CA).

## FUNDING

This work was funded by a grant from the National Institute of Health (R01 AG051810) to L.R.L.

## AUTHOR CONTRIBUTIONS

S.Q.W conceived the experiments, generated strains, performed imaging, lifespan, heat stress, and data analyses, and wrote the manuscript. C.J.R performed lifespan and heat stress analyses, and provided feedback on the manuscript. L.R.L provided conceptual feedback, performed lifespan analyses and edited the manuscript.

## ACKNOWLEDGEMENTS

We are grateful to Dr. Anita V. Kumar, Dr. Joslyn Mills, and Dennis M. Bonal (Brown University, RI) for their valuable insights and suggestions towards the study and manuscript preparation. We would also like to extend gratitude to Dr. Andrew Dillin (HHMI/UC Berkeley, CA) for providing constructs pLp17 and pLP19. We thank the Caenorhabditis Genetics Center (U. Minnesota, P40 OD010440) for providing transgenic and mutant strains.

## COI STATEMENT

The authors declare that they have no competing interests.

---

**Supplemental Figure 1.**
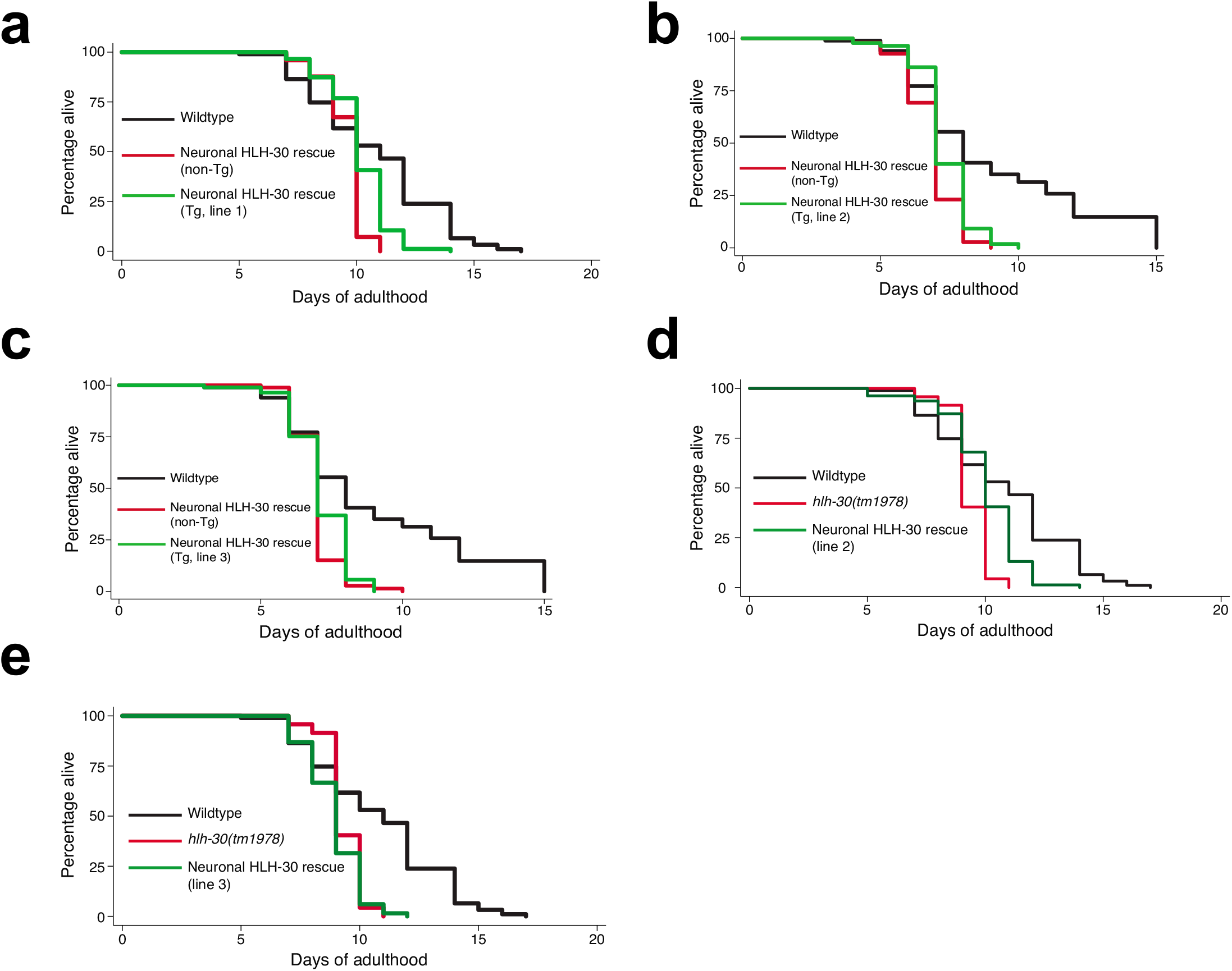
Neuronal HLH-30/TFEB does not regulate normal lifespan. Lifespan analyses of *hlh-30(tm1978)* mutants (as non-transgenic (non-Tg) siblings and *hlh-30(tm1978)*) rescued with **(a to c)** extrachromosomal (Tg, transgenic) and **(d and e)** integrated arrays of HLH-30/TFEB in neurons on OP50 at 25°C. Animals were developed at 20°C and shifted to 25°C on OP50 from day 1 of adulthood. Data are representatives of **(a)** 4, (**b** and **c)** single, and (**d** and **e)** 3 independent replicates, and comparisons were made by Mantel-Cox log-rank. Further details about lifespan analyses are provided in Supplemental Table 1.

**Supplemental Figure 2.**
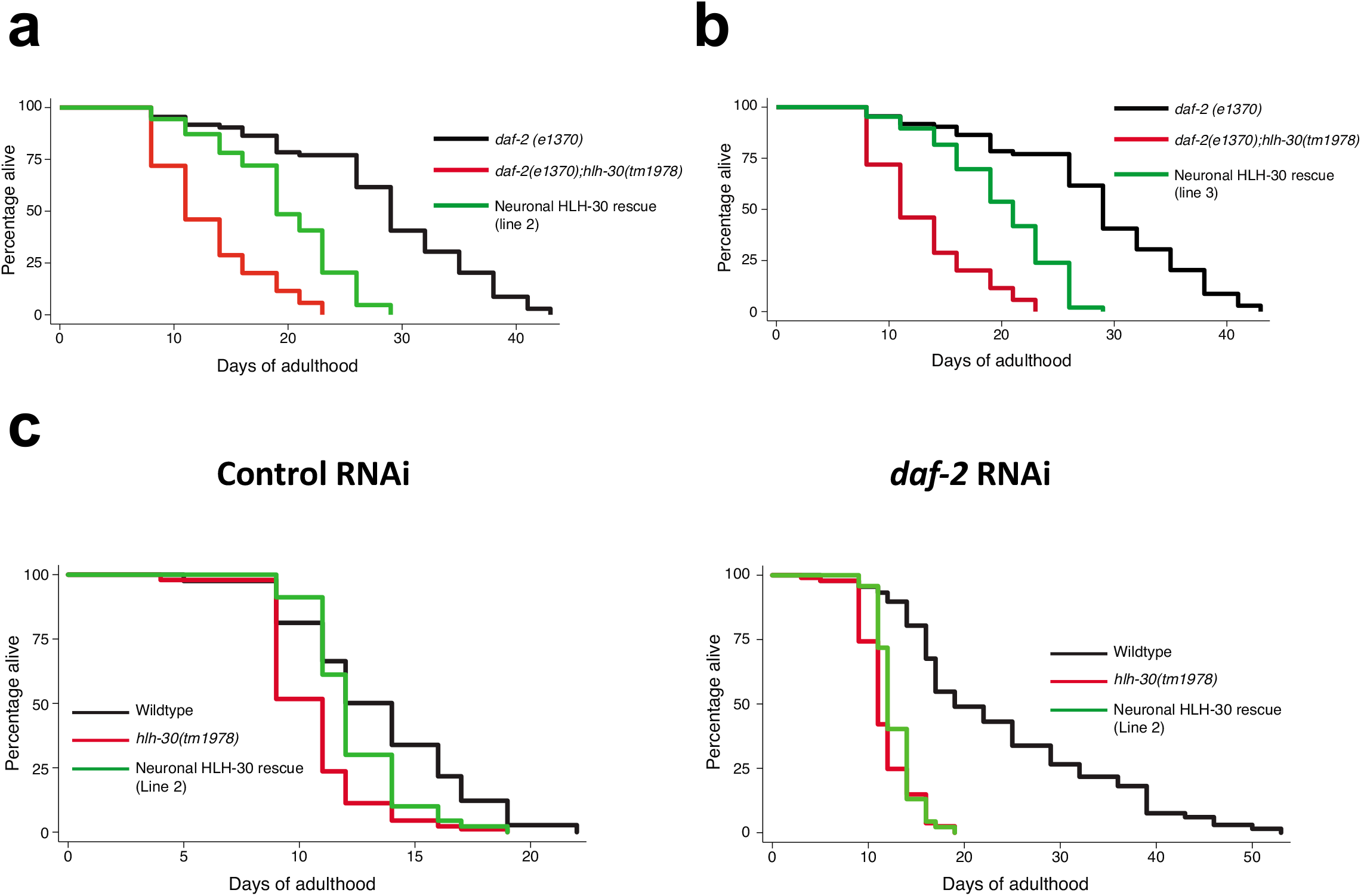
Neuronal HLH-30/TFEB regulates longevity dependently of *daf-16*. **(a and b)** Lifespan analyses of daf-2(e1370), daf-2(e1370);hlh-30(tm1978), and neuronal HLH-30/TFEB rescued animals fed OP50 at 25°C. Animals were developed at 20°C and shifted to 25°C on OP50 from day 1 of adulthood. Data are representatives of 3 independent replicates, and comparisons were made by Mantel-Cox log-rank. Further details about lifespan analyses are provided in Supplemental Table 2. **(c)** Lifespan analyses of of wildtype, *hlh-30(tm1978)*, and neuronal HLH-30/TFEB rescued animals fed control RNAi (*L4440*) or RNAi against *daf-2* at 25°C. Note that neuronal HLH-30/TFEB rescued animals from the only transgenic line (line 2) exhibiting lifespan rescue in comparison to *hlh-30(tm1978)* mutants were used for analyses here **(Refer to** Supplemental Figure 1d **and Table 1)**. Animals developed at 20°C on OP50 were transferred on day 1 of adulthood onto bacteria expressing either control dsRNA (*L4440*) or dsRNA against *daf-2* and grown at 25°C. Data are representatives of 2 independent replicates and comparisons were made by Mantel-Cox log-rank. Further details about lifespan analyses are provided in Supplemental Table 4.

**Supplemental Figure 3.**
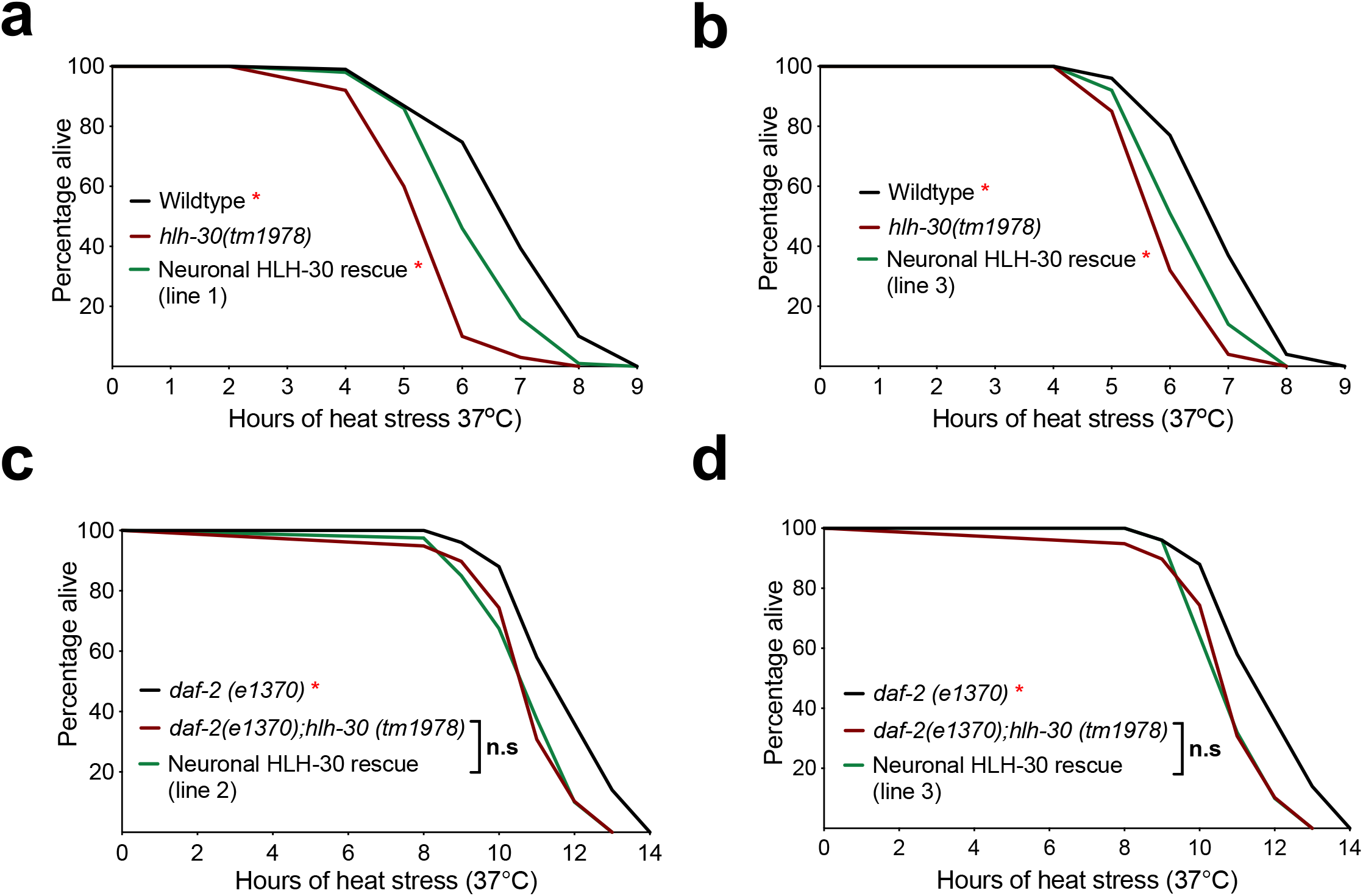
Neuronal HLH-30/TFEB mediates thermoresistance in normal but not longevity-promoting conditions. Survival analyses of neuronal HLH-30/TFEB rescued animals in comparison with their **(a and b)** wildtype and *hlh-30(tm1978)* and **(c and d)** *daf-2(e1370* and *daf-2(e1370);hlh-30(tm1978)* counterparts at 37°C heat stress. Animals were developed at 20°C and shifted to heat stress at 37°C on day 1 of adulthood. Data are representatives of (**a** and **b)** single experiments and **(c and d)** 2 independent replicates and comparisons were made by Mantel-Cox log-rank (*n* = 90-100/strain; n.s, *p*≥0.05; *, *p*<0.05; in comparison to *hlh-30(tm1978)* or *daf-2(e1370);hlh-30(tm1978))*.

**Supplemental Figure 4.**
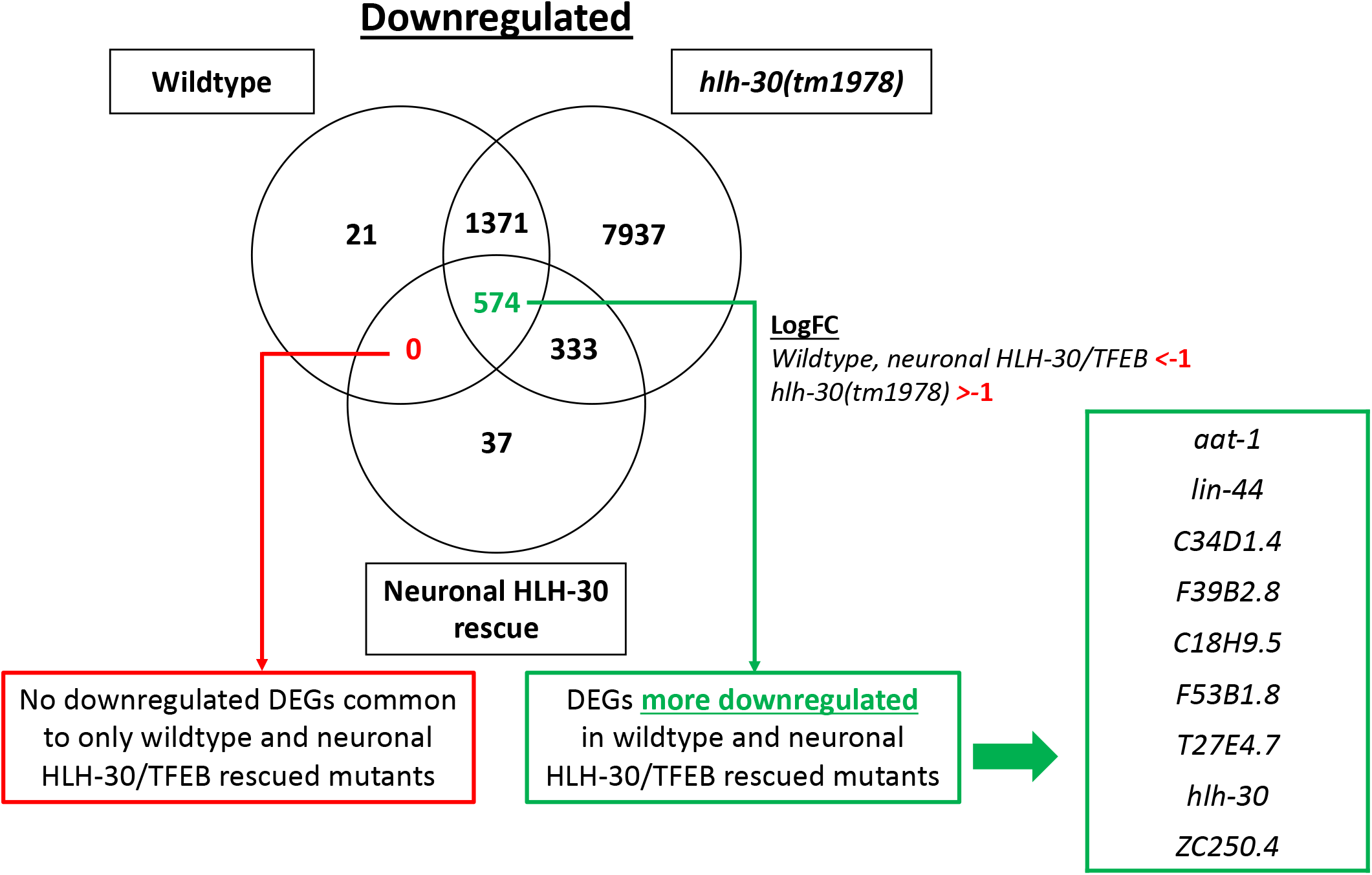
Downregulated differentially expressed genes (DEGs) in wildtype and neuronal HLH-30/TFEB animals in comparison to *hlh-30(tm1978)* mutants. Pairwise comparisons of RNAseq profiles of day 1 animals from 20°C (control conditions) and 3 hrs heat stress at 37°C identified genotype-specific differentially expressed genes (DEGs) downregulated with heat stress, which were overlapped between genotypes to extract significant hits (adjusted *p*<0.05) common to or more downregulated (Log_2_ fold change (LogFC) thresholds applied as indicated) in wildtype and neuronal HLH-30/TFEB rescued animals than *hlh-30(tm1978)* mutants. Animals from 4 independent replicates were developed at 20°C to day 1 of adulthood and harvested for RNA after 3 hrs of further growth at 20°C or 37°C.

**Supplemental Figure 5.**
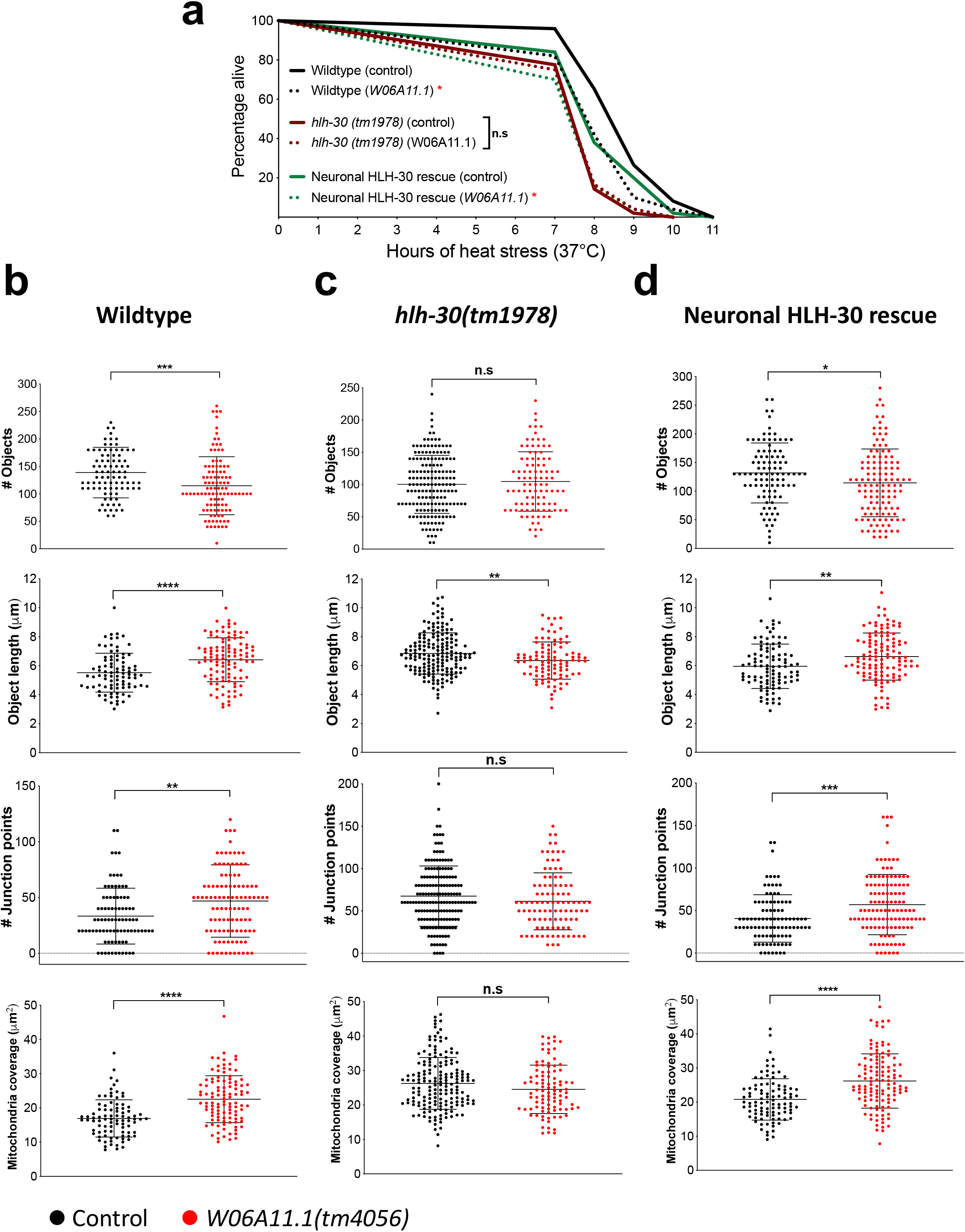
Neuronal HLH-30/TFEB mediates thermoresistance through W06A11.1-dependent mitochondrial fragmentation in muscles. **(a)** Survival analyses of wildtype, *hlh-30(tm1978)*, and neuronal HLH-30/TFEB rescued animals fed control RNAi (*L4440*) or RNAi against *W06A11.1* at 37°C heat stress. Animals developed at 20°C to day 1 of adulthood on OP50 were transferred onto bacteria expressing either control dsRNA (*L4440*) or dsRNA against *W06A11.1*, grown at 25°C for 48 hr, and exposed to 37°C heat stress. Data is representative of 2 independent replicates (*n* = 83 – 99/RNAi/strain) and comparisons were made by Mantel-Cox log-rank (n.s, *p*≥0.05; *, *p*<0.05; in comparison to control RNAi). **(b to d)** Analysis of mitochondrial connectivity in the absence and presence of *W06A11.1(tm4056)* loss of function after 37°C heat stress for 3 hrs in **(b)** wildtype, **(c)** *hlh-30(tm1978)* loss of function mutants, and **(d)** neuronal HLH-30/TFEB rescued animals. Animals were developed at 20°C to day 1 of adulthood and exposed to 37°C heat stress for 3 hrs. Data are representatives of 3 independent replicates (per strain; *n =* 30, number of ROIs = 88 – 171). Comparisons were made by either Mann-Whitney or student’s unpaired *t-*test accordingly to the normality of data distribution for each analyzed mitochondrial feature (presented as mean ± S.D; n.s, *p*≥0.05; *, *p*<0.05; **, *p*<0.01; ***, *p*<0.001; ****, *p*<0.0001).

**Supplemental Figure 6.**
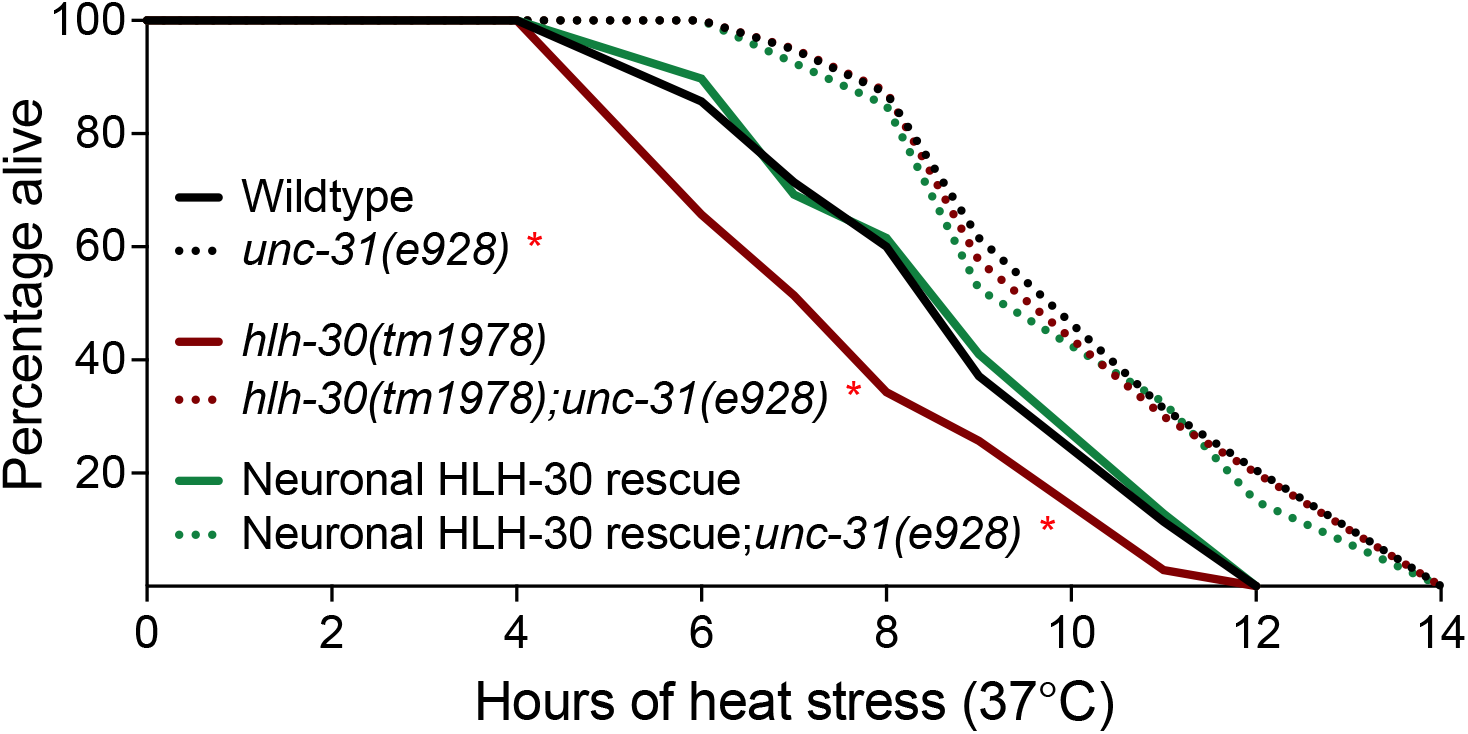
Neuronal HLH-30/TFEB does not mediate thermoresistance through dense core vesicle (DCV) release. Survival analyses of wildtype, *hlh-30(tm1978)*, and neuronal HLH-30/TFEB rescued animals in the absence and presence of *unc-31(e928)* loss of function at 37°C heat stress. Animals were developed at 20°C to day 1 of adulthood and exposed to 37°C heat stress. Data is representative of 3 – 4 independent replicates and comparisons were made by Mantel-Cox log-rank (*n* = 113 – 169/strain; *, *p*<0.05; comparisons of *unc-31(e928)* to control per genotype).

**Supplemental Figure 7.**
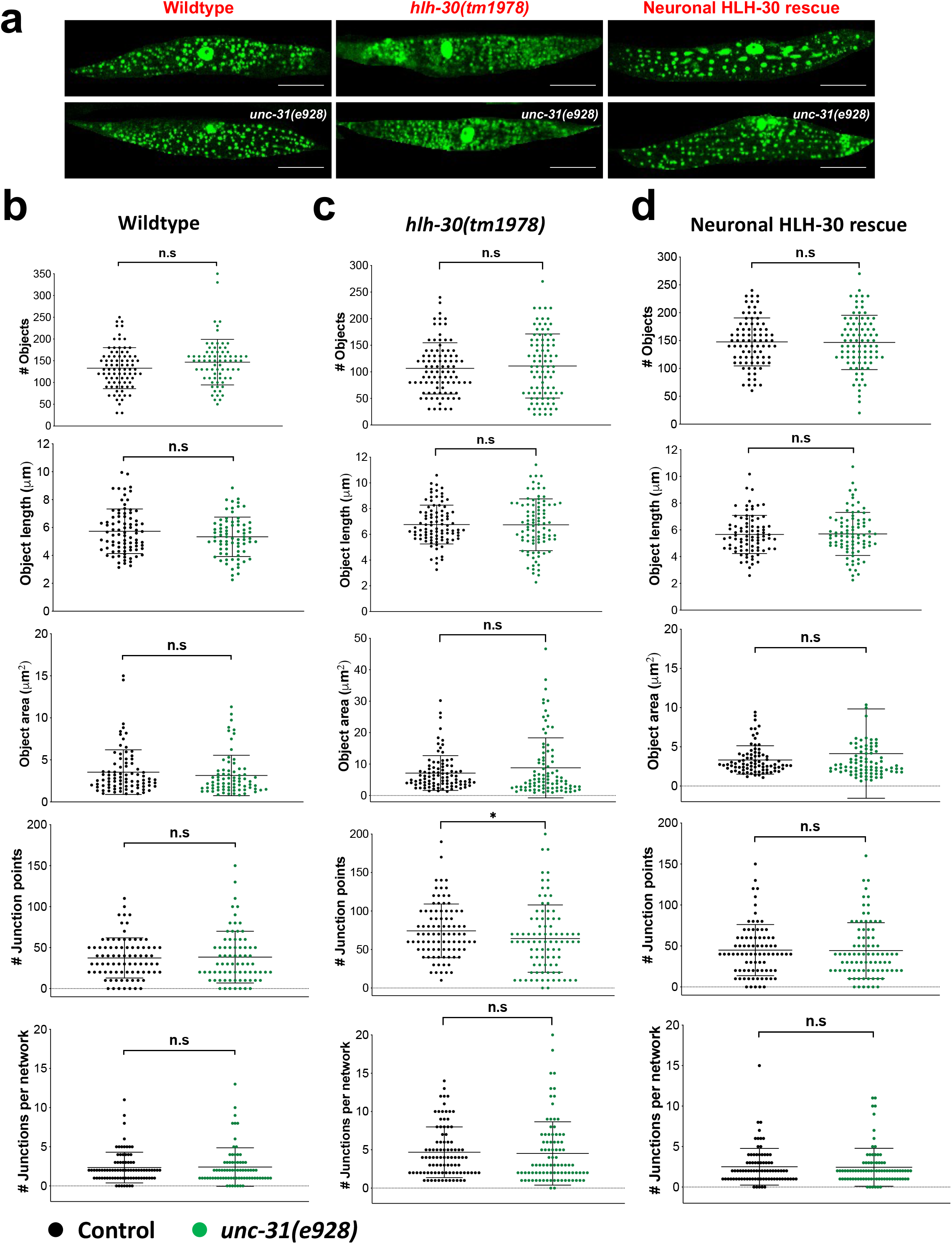
Heat stress-induced mitochondrial fragmentation was not dependent on signaling by dense core vesicle (DCV) release. **(a)** Representative images of muscle mitochondrial morphology with the body wall muscle mitochondrial reporter (Mito::GFP) in wildtype, *hlh-30(tm1978)*, and neuronal HLH-30/TFEB rescued animals in the **(top panel)** absence and **(bottom panel)** presence of *unc-31(e928)* loss of function after 37°C heat stress for 3 hrs. **(b to d)** Analysis of mitochondrial connectivity in the absence and presence of *unc-31(e928)* loss of function in **(b)** wildtype, **(c)** *hlh-30(tm1978)*, and **(d)** neuronal HLH-30/TFEB rescued animals after 37°C heat stress for 3 hrs. Animals were developed at 20°C to day 1 of adulthood and exposed to 37°C heat stress for 3 hrs. Data are representatives of 3 independent replicates (per strain; *n* = 30, number of ROIs= 75 – 92) and comparisons were made by either Mann-Whitney or student’s unpaired *t-*test accordingly to the normality of data distribution for each analyzed mitochondrial feature (presented as mean ± S.D; n.s, *p*≥0.05; *, *p*<0.05; **, *p*<0.01; ***, *p*<0.001; ****, *p*<0.0001). Scale bars = 20 μM.

**Supplemental Figure 8.**
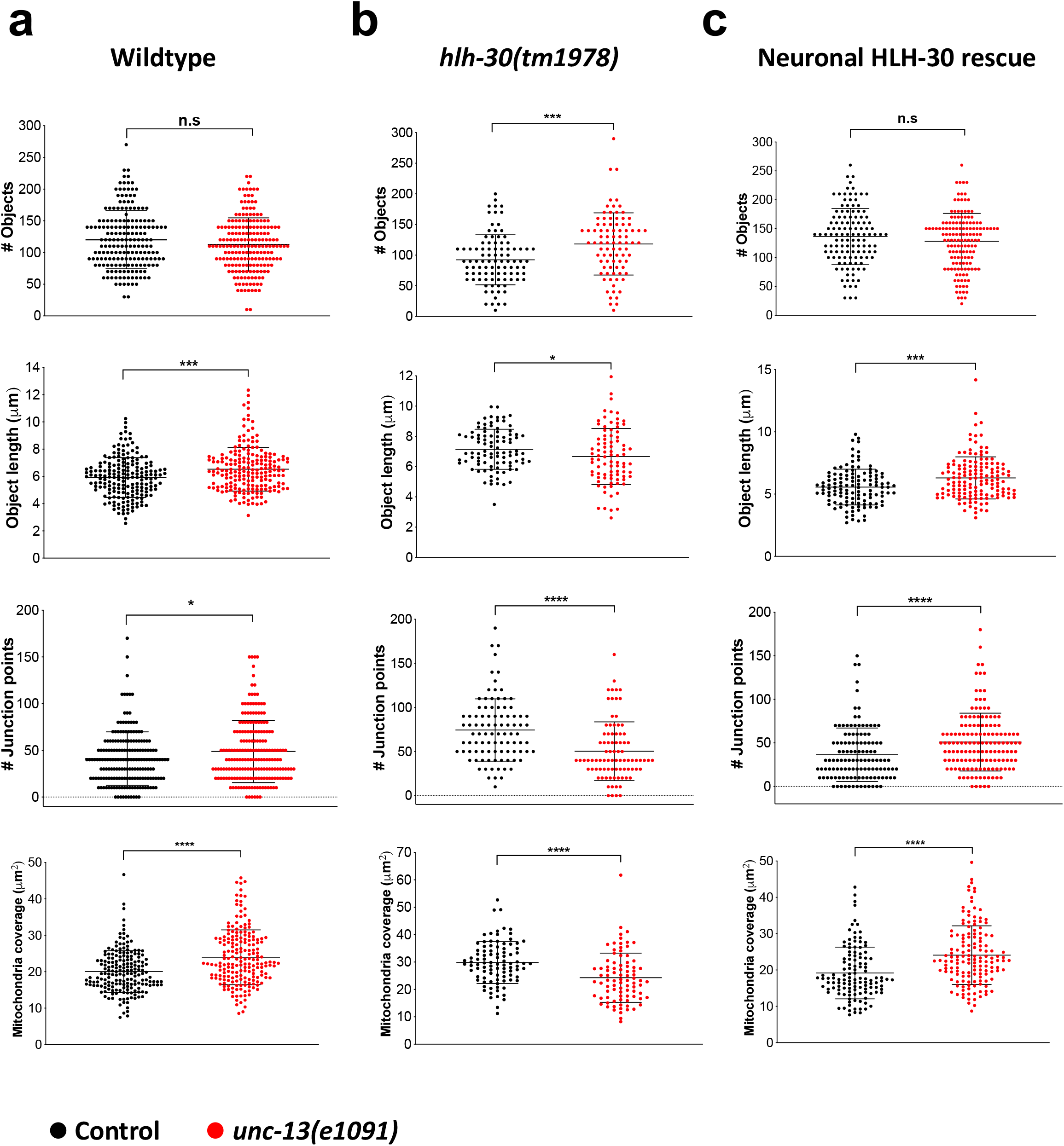
Defective neurotransmission increases heat stress-induced mitochondria fragmentation in *hlh-30(tm1978)* mutants. Analysis of mitochondrial connectivity in the absence and presence of *unc-13(e1091)* loss of function in **(a)** wildtype, **(b)** *hlh-30(tm1978)*, and **(c)** neuronal HLH-30/TFEB rescued animals after 37°C heat stress for 3 hrs. Animals were developed at 20°C to day 1 of adulthood and exposed to 37°C heat stress for 3 hrs. Data are representatives of 3 - 5 independent replicates (per strain; *n* = 30 - 50; number of ROIs = 87 – 196) and comparisons were made by either Mann-Whitney or student’s unpaired *t-*test accordingly to the normality of data distribution for each analyzed mitochondrial feature (presented as mean ± S.D; n.s, *p*≥0.05; *, *p*<0.05; **, *p*<0.01; ***, *p*<0.001; ****, *p*<0.0001). Scale bars = 20 μM.

**Supplemental Table 1.** Lifespan analyses of neuronal HLH-30/TFEB rescued animals in *hlh-30(tm1878)* background. Animals were raised at 20°C and grown at 25°C on OP50. Where indicated (*), low number of events observed (EO, <50) were due largely due to censoring of animals which were more susceptible to internal progeny hatching at 25°C. Statistical analyses: Mantel-Cox log-rank; Mean LS: Mean Lifespan. Refer to Supplemental Table 5 for additional strain information.

| N2 |  | *hlh-30(tm1978)* |  |  |  | Neuronal HLH-30 rescue strains |  |  |  |  |  |
| --- | --- | --- | --- | --- | --- | --- | --- | --- | --- | --- | --- |
| Mean LS | EO | Strain | Mean LS | EO | % Difference to N2 | Strain | Mean LS | EO | % Difference to *hlh-30(tm1978)* | p Value | Figure # |
| 9.02 | 68 | LRL81 (Non Tg) | 6.9 | 83 | -23.53 | LRL81 (Tg) | 7.34 | 68 | 6.42% | 0.0012 |  |
| 12.99 | 47 | LRL81 (Non Tg) | 9.57 | 74 | -26.33% | LRL81 (Tg) | 10.16 | 73 | 6.17% | 0.0070 |  |
| 10.79 | 93 | LRL81 (Non Tg) | 9.58 | 98 | -11.21% | LRL81 (Tg) | 10.14 | 86 | 5.85% | < 0.0001 | S1a |
| 9.48 | 74 | LRL81 (Non Tg) | 8.44 | 92 | -0.10% | LRL81 (Tg) | 8.96 | 71 | 6.16% | 0.0086 |  |
| 9.02 | 57 | LRL82 (Non Tg) | 6.86 | 88 | -23.95% | LRL82 (Tg) | 7.32 | 65 | 6.70% | 0.0027 | S1b |
| 9.02 | 57 | LRL83 (Non Tg) | 6.94 | 84 | -23.05% | LRL83 (Tg) | 7.12 | 71 | 2.65% | 0.0736 | S1c |
| 12.99 | 47 * | LRL31 | 9.46 | 59 | -27.17% | LRL146 (Line 1) | 9.71 | 77 | 2.64% | 0.3818 |  |
| 10.79 | 93 | LRL31 | 9.32 | 93 | -13.62% | LRL146 (Line 1) | 9.29 | 89 | -0.32% | 0.4978 | 1b |
| 9.48 | 74 | LRL31 | 7.7 | 88 | -18.78% | LRL146 (Line 1) | 7.91 | 86 | 2.73% | 0.2631 |  |
| 12.99 | 47 * | LRL31 | 9.46 | 59 | -27.17% | LRL147 (Line 2) | 10.36 | 62 | 9.51% | < 0.0001 |  |
| 10.79 | 93 | LRL31 | 9.32 | 93 | -13.62% | LRL147 (Line 2) | 9.98 | 77 | 7.02% | < 0.0001 | S1d |
| 9.48 | 74 | LRL31 | 7.7 | 88 | -18.78% | LRL147 (Line 2) | 8.7 | 75 | 12.99% | < 0.0001 |  |
| 12.99 | 47 * | LRL31 | 9.46 | 59 | -27.17% | LRL148 (Line 3) | 9.49 | 48 * | 0.32% | 0.9957 |  |
| 10.79 | 93 | LRL31 | 9.32 | 93 | -13.62% | LRL148 (Line 3) | 8.93 | 82 | -4.18% | 0.0424 | S1e |
| 9.48 | 74 | LRL31 | 7.7 | 88 | -18.78% | LRL148 (Line 3) | 7.56 | 75 | -1.82% | 0.5542 |  |

**Supplemental Table 2.** Lifespan analyses of neuronal *hlh-30* rescued animals in *daf-2(e1370);hlh-30(tm1978)* background. Animals were raised at 20°C and grown at 25°C on OP50. Where indicated (*), low number of events observed (EO, <50) were due largely due to censoring of animals which were more susceptible to internal progeny hatching at 25°C. Statistical analyses: Mantel-Cox log-rank; Mean LS: Mean Lifespan. Refer to Supplemental Table 5 for additional strain information.

| *daf-2(e1370)III* |  | *daf-2(e1379)III;*<br>*hlh-30(tm1978)IV* |  |  | Neuronal HLH-30 rescue strains |  |  |  |  |  |
| --- | --- | --- | --- | --- | --- | --- | --- | --- | --- | --- |
| Mean LS | EO | Mean LS | EO | % Difference to *daf-2(e1370)III* | Strain | Mean LS | EO | % Difference to *daf-2(e1370)III;*<br>*hlh-30(tm1978)* | p Value | Figure # |
| 31.52 | 95 | 15.28 | 69 | -51.52% | **LRL167**<br>(Line 1) | 18.8 | 70 | 23.04% | < 0.0001 |  |
| 28.35 | 72 | 13.06 | 50 | -53.93% | **LRL167**<br>(Line 1) | 22.07 | 70 | 68.98% | < 0.0001 | 1c |
| 33.67 | 80 | 13.26 | 61 | -60.62% | **LRL167**<br>(Line 1) | 20.91 | 73 | 57.69% | < 0.0001 |  |
| 31.52 | 95 | 15.28 | 69 | -51.52% | **LRL168**<br>(Line 2) | 19.35 | 84 | 26.64% | < 0.0001 |  |
| 28.35 | 72 | 13.06 | 50 | -53.93% | **LRL168**<br>(Line 2) | 19.71 | 66 | 50.92% | < 0.0001 | S2a |
| 33.67 | 80 | 13.26 | 61 | -60.62% | **LRL168**<br>(Line 2) | 23.08 | 65 | 74.06% | < 0.0001 |  |
| 31.52 | 95 | 15.28 | 69 | -51.52% | **LRL169**<br>(Line 3) | 18.07 | 61 | 18.26% | 0.0009 |  |
| 28.35 | 72 | 13.06 | 50 | -53.93% | **LRL169**<br>(Line 3) | 19.69 | 52 | 52.35% | < 0.0001 | S2b |
| 33.67 | 80 | 13.26 | 61 | -60.62% | **LRL169**<br>(Line 3) | 19.16 | 39 * | 44.40% | < 0.0001 |  |

**Supplemental Table 3.** Lifespan analyses of neuronal *hlh-30* rescued animals in *daf-2(e1370);hlh-30(tm1978)* background on *daf-16* RNAi. Animals developed at 20°C on OP50 were transferred on day 1 of adulthood onto bacteria expressing either control dsRNA (*L4440*) or dsRNA against *daf-16* and grown at 25°C. Where indicated (*), low number of events observed (EO, <50) were largely due to censoring of animals which were more susceptible to internal progeny hatching at 25°C. Statistical analyses: Mantel-Cox log-rank; Mean LS: Mean Lifespan. Refer to Supplemental Table 5 for additional strain information.

|  | Control RNAi |  |  |  | *daf-16* RNAi |  |  |  |  |
| --- | --- | --- | --- | --- | --- | --- | --- | --- | --- |
| Strain | Mean LS | EO | % Difference to *daf-2(e1370)III* | *p* Value | Mean LS | EO | % Difference to *daf-2(e1370)III* | *p* Value | Figure # |
| *daf-2(e1370)III* | 22.96 | 67 |  |  | 21.87 | 93 |  |  |  |
| *daf-2(e1379)III; hlh-30(tm1978)IV* | 12.47 | 36 * | -45.69 | < 0.0001 | 14.58 | 90 | -33.33 | < 0.0001 |  |
| Neuronal HLH-30 rescue (LRL167) | 18.18 | 43 * | -20.82 | 0.0133 | 14.66 | 83 | -32.97 | < 0.0001 |  |
| *daf-2(e1370)III* | 31.24 | 82 |  |  | 20.16 | 74 |  |  | 1d |
| *daf-2(e1379)III; hlh-30(tm1978)IV* | 15.05 | 37 * | -51.82 | < 0.0001 | 15.02 | 61 | -25.5 | < 0.0001 |  |
| Neuronal HLH-30 rescue (LRL167) | 25.19 | 43 * | -19.37 | 0.0019 | 14.5 | 46 * | -28.08 | < 0.0001 |  |
| *daf-2(e1370)III* | 27.27 | 61 |  |  | 15.90 | 61 |  |  |  |
| *daf-2(e1379)III; hlh-30(tm1978)IV* | 13.09 | 20 * | -52 | < 0.0001 | 12.86 | 60 | -19.11 | < 0.0001 |  |
| Neuronal HLH-30 rescue (LRL167) | 18.54 | 40 * | -32.01 | < 0.0001 | 12.92 | 57 | -18.74 | < 0.0001 |  |

**Supplemental Table 4.** Lifespan analyses of neuronal *hlh-30* rescued animals in *hlh-30(tm1878)* background on *daf-2* RNAi. Animals raised at 20°C on OP50 were transferred on day 1 of adulthood onto bacteria expressing either control dsRNA (*L4440*) or dsRNA against *daf-2* and grown at 25°C. Statistical analyses: Mantel-Cox log-rank; Mean LS: Mean Lifespan; EO: Events Observed. Refer to Supplemental Table 5 for additional strain information. Note that neuronal HLH-30/TFEB rescued animals from the only transgenic line (line 2) which exhibited partial lifespan rescue in comparison to *hlh-30(tm1978)* mutants were used for analyses here **(Refer to** Supplemental Figure 1d **and Table 1)**.

|  | Control RNAi |  |  |  | *daf-2* RNAi |  |  |  |  |
| --- | --- | --- | --- | --- | --- | --- | --- | --- | --- |
| Strain | Mean LS | EO | % Difference to N2 | *p* Value | Mean LS | EO | % Difference to N2 | *p* Value | Figure # |
| N2 | 12.29 | 62 |  |  | 19.52 | 51 |  |  |  |
| *hlh-30(tm1978)* | 9.58 | 61 | -22.05 | < 0.0001 | 11.84 | 55 | -39.34 | < 0.0001 |  |
| Neuronal HLH-30 rescue (LRL147; Line 2) | 12.21 | 78 | -0.65 | 0.5442 | 14.65 | 74 | -24.95 | < 0.0001 |  |
| N2 | 13.41 | 74 |  |  | 23.90 | 82 |  |  | S2c |
| *hlh-30(tm1978)* | 10.53 | 90 | -21.52 | < 0.0001 | 11.68 | 81 | -51.14 | < 0.0001 |  |
| Neuronal HLH-30 rescue (LRL147; Line 2) | 12.33 | 90 | -8.12 | 0.0027 | 12.79 | 92 | -46.50 | < 0.0001 |  |

**Supplemental Table 5.**
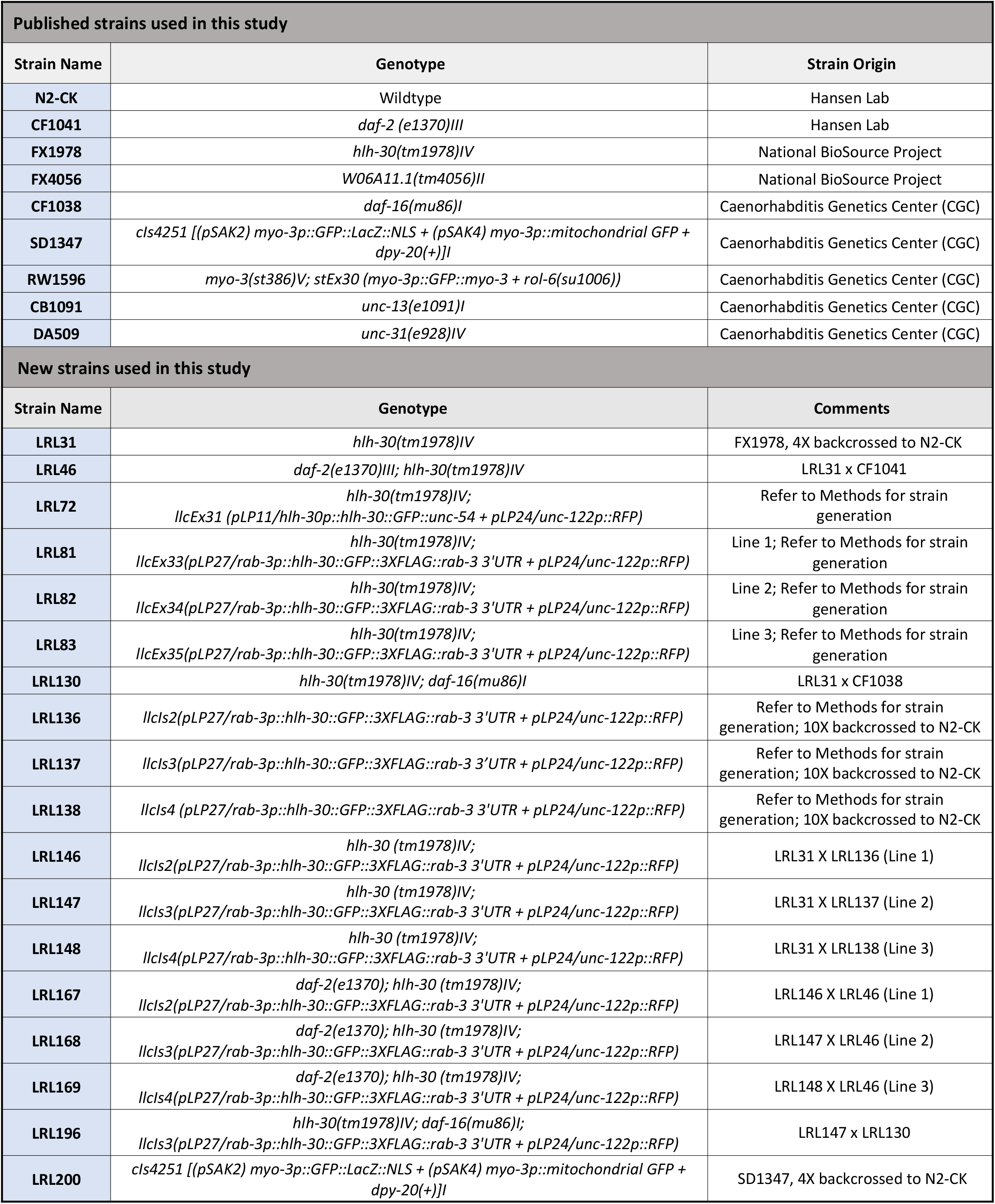
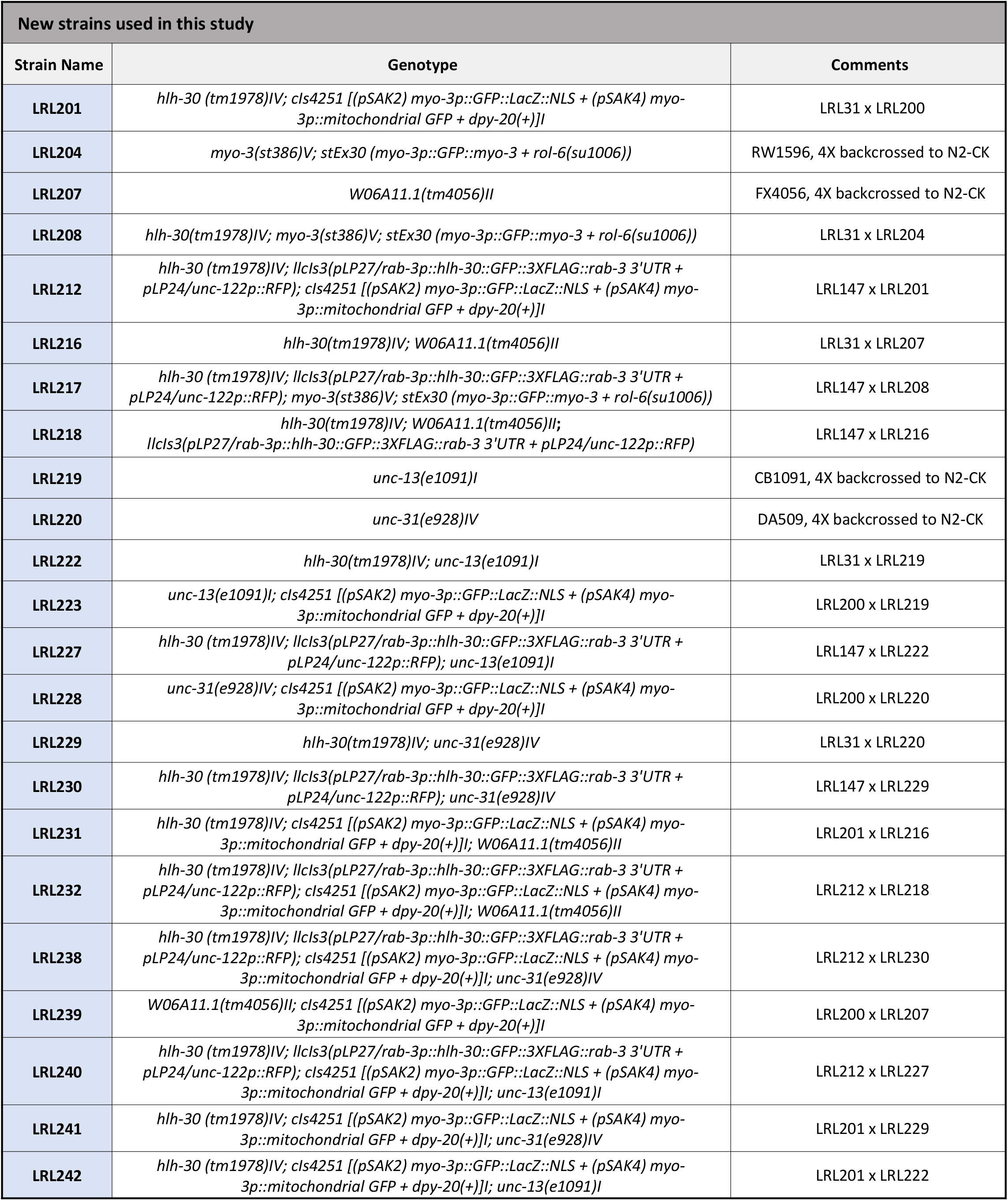
Strains used in the study.

**Supplemental Table 6.**
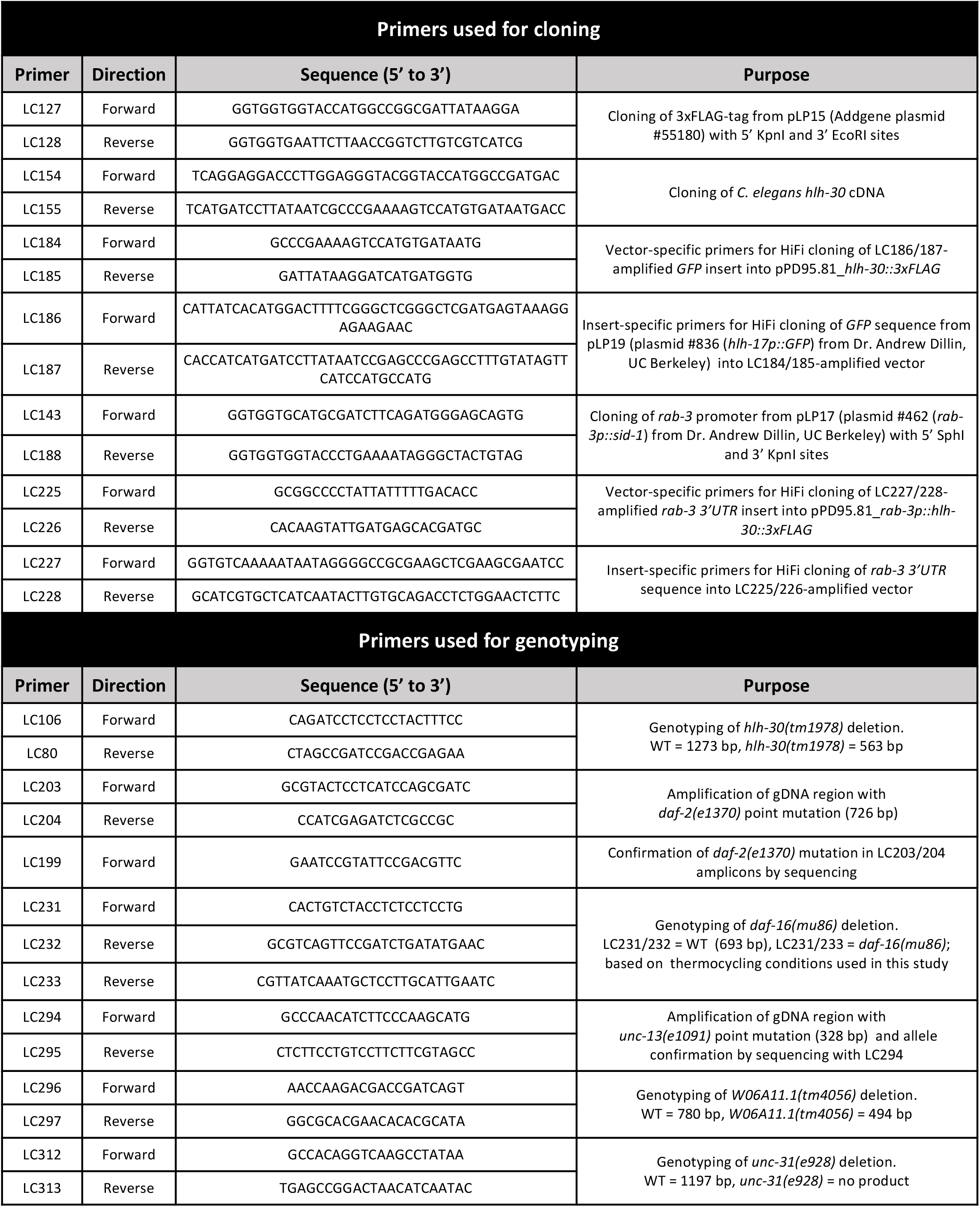
Primers used in this study.

## Notes

### Competing Interest Statement

The authors have declared no competing interest.

